# Higher frequency of transition mutation over transversion mutation in genomes: evidence from the binding energy calculation of base pairs using DFT

**DOI:** 10.1101/2024.11.28.625875

**Authors:** Nishita Deka, Nand Kishor Gour, Pradeep Pant, Siddhartha Shankar Satapathy, Nasimul Hoda, Ramesh Chandra Deka, Suvendra Kumar Ray

**Affiliations:** Department of Molecular Biology and Biotechnology, Tezpur University, Tezpur, 784028 Assam, India; Department of Chemical Sciences, Tezpur University, Tezpur, 784028 Assam, India; Department of Biotechnology, Bennett University, Greater Noida 201310, Uttar Pradesh; Department of Computer Science & Engineering, Tezpur University, Tezpur, 784028 Assam, India; Department of Chemistry, Jamia Millia Islamia, Jamia Nagar,110025, New Dehli; Centre for Multidisciplinary Research, Tezpur University, Tezpur, 784028 Assam, India; Centre for Bioinformatics & Computational Biology, Tezpur University, Tezpur, 784028 Assam, India

**Keywords:** Base substitution mutation, Base pairs, Transition, Transversion, DFT, BSSE, Binding energy, Base tautomer, Base conformers

## Abstract

Base substitution mutations such as transition (*ti*) and transversion (*tv*) in organisms are major driving force in molecular evolution. In this study, different possible types of base pairing that can cause *ti* and *tv* were investigated using the density functional theory (DFT) method. The chemical structures of bases as well as base pairs were optimized using B3LYP hybrid functional along with 6-31G(d,p) basis set. We performed single point energy calculation of all optimized species using the same functional but combined with higher diffuse and polarized basis set i.e. 6-311++G(d,p) to get more refined energy of all species. The binding energy of various base pairs was calculated considering basis set superposition error (BSSE) as well as without BSSE. The binding energy of the base pairs leading to *ti* was found to be more stable than that of the base pairs leading to *tv*. This was interesting considering the observations in organisms that *tis* are more frequent than *tvs*. Among the base pairs leading to the same *ti*, G(*keto*): T (*enol)* base pair was found to be more stable than A(*imino*):C(*amino)* base pair. This theoretical study of binding energy of different base pairs using the DFT method has provided additional evidences in support to the biological observations of a higher transition rate than transversion in genomes.

## 1. Introduction

Base substitution mutation in DNA is the most common mechanism of genome variation in organisms. These mutations have been classified into two types such as transitions (*ti*) where, a purine (or a pyrimidine) base is substituted by another purine (or pyrimidine) base, and transversions, where a purine base is substituted by a pyrimidine base, or the *vice versa* (Sankoff et al., 1976).

Considering that each base in DNA can be replaced by any of the other three bases, there are a total of twelve possible substitutions, of which eight are transversions and four are transitions (Sankoff et al., 1976). The transitions are C→T, T→C, A→G, and G→A which result from purine-pyrimidine (Pu:Py) base pairing. In transversions, purine-purine (Pu-Pu) or pyrimidine-pyrimidine (Py-Py) base pairing (Leonard et al.,1994) leading to the following transversions: A→T, A→C, T→A, T→G, G→T, G→C, C→A, and C→G. Although, theoretically transversions should occur more frequently, transition mutations are observed to occur two to three times more frequent than transversions in genomes (Vogel & Kopun, 1977;(Sen et al., 2021); Aziz et al., 2022). This disparity has been mainly attributed to the distortion of the DNA helix geometry caused by Pu-Pu and Py-Py pairs responsible for *tv* compared to Pu-Py pair responsible for *ti* in genomes (Topal & Fresco, 1976). Further, among the different *ti*s, the frequencies of C→T (or G→A) is higher than that of T→C (or A→G) (Kreutzer & Essigmann, 1998). There is also difference among the eight *tv*s: G→T (or C→A) usually occurs at a higher frequency than that of the other *tv*s (Hollstein et al., 1991). Base tautomer, such as amino/imino and keto/enol forms, and base conformers, like anti/syn conformations, play a significant role in forming base pairs that lead to these substitutions (Table 1). Tautomerization primarily contributes to transitions, while the tautomer along with syn conformers facilitate transversions (Topal & Fresco, 1976). Though factors such as DNA geometry, DNA error repair enzymes, damage to DNA bases, RNA secondary structure and codon degeneracy have been implemented towards *ti*s and *tv*s in genomes, there is no report in the literature regarding the base pair stability in DNA with their contribution towards these mutation types. By estimating the binding energies of these base pairs (Table 1), their stability can be determined — the lower the binding energy, the more stable the pairing.

**Table 1.** Base pairs that cause different types of base substitution mutation are shown.

| 12 substitutions | Base pairs |
| --- | --- |
| A→C | A: G |
| A→G | G: T or A:C |
| A→T | A: A |
| C→A | G: A |
| C→G | G: G |
| C→T | G: T or A:C |
| G→A | G: T or A:C |
| G→C | G: G |
| G→T | A: G |
| T→A | A: A |
| T→C | A:C or G: T |
| T→G | A: G |

The binding energy of DNA base pairs can be estimated through experimental methods and computational analysis using quantum chemical approaches. The binding energies of the A: T and G:C base pairs have been empirically estimated with values of - 13.00 kcal/mol and -21.00 kcal/mol, respectively (Gould & Kollman, 1994; Yanson et al., 1979). With advancements in quantum chemical studies, researchers are now able to estimate the binding energies of base pairs that are difficult to measure experimentally. For example, rare tautomer and base conformers are very short-lived, making it challenging to experimentally determine their stability. Quantum chemical approaches have proven to be promising, as they can mimic experimental results. Several researchers have explored the binding energy between DNA base pairs using computational methods, yielding results similar to experimental data. For example, the binding energy of the A: T base pair has been reported to range from -9.00 to -17.00 kcal/mol (Souri & Mohammadi, 2017; Masoodi et al., 2019), while the binding energy of the G:C base pair has been found to range from -22.00 to - 31.00 kcal/mol (Asensio et al., 2003; Acosta-Silva et al., 2010; Thoa & Hue, 2018). Masoodi et al. (2018) investigated the binding energies of A _*imino*_: C _*amino*_ and A _*amino*_: C _*imino*_ base pairs using the DFT method, reporting values of -21.00 kcal/mol and -14.00 kcal/mol, respectively. However, base pairs formed by tautomerization as well as base conformational changes (anti/syn), which contribute to base substitution mutations, have not been extensively studied previously. In this work, we have explored binding energy of different base pairs that have undergone tautomerization, conformational changes, or both. The base pairs we considered (that cause mutation) are based on the hypothesis of Topal and Fresco (1976). Unlike earlier studies, which typically excluded the sugar-phosphate backbone and focused solely on the bases, we included the nucleotide pair, complete with its sugar-phosphate backbone, to more accurately represent the base conformers. We have observed that base pairs leading to tis have lower binding energy than that of the base pairs leading to tvs. This approach has allowed us to provide another explanation for the higher transition-to-transversion *(ti/tv)* ratio in genomes, supported by quantum chemical analysis using the DFT method.

## 2. Materials and Computational Details

All the base structures were drawn using Gauss View 6, starting from nucleotide templates provided by the software and applying the desired modifications. We assumed that the nucleotide templates were accurate and proceeded accordingly. After that, the hydrogen bond is added between the respective pairs. All DFT investigations were executed using the GAUSSIAN 16 software package.(Frisch et al., 2016). Geometry optimizations of single bases and base pairs were conducted using B3LYP functional (Lee et al., 1988) along with 6-31G (d, p) basis set (Muzomwe et al., 2012),(Bouzzine et al., 2005),(Tirado-Rives & Jorgensen, 2008) B3LYP functional incorporates Becke’s three-parameter hybrid exchange functional and the Lee-Yang–Parr correlation functional. Vibrational frequency calculations are further performed at the same level of theory to get different modes of vibrational frequency as well as verify all the stable minima to have positive frequencies. Single point energy calculation of all optimized species were further performed at the same functional but using higher diffuse and polarized basis set i.e. 6-311++G (d,p) to get more refined energy of all species (Andersson & Uvdal, 2005), (Appell et al., 2005), (Schnupf et al., 2007). Corrections for basis set superposition error (BSSE) and zero-point vibrational energies (ZPE) for binding energies were taken into account by using the counterpoise (CP) method (Gutowski et al., 1993),(Tao & Pan, 1991),(Novoa et al., 1994)

The binding energy (ΔE) was determined as the difference in energy between the dimer and its isolated monomers. All calculations included counterpoise corrections for BSSE and adjustments for zero-point vibrational energies.

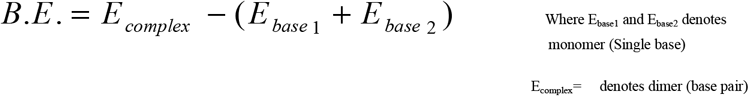

## 3. Results

### Binding energy of base pairs leading to transition is more stable than the base pairs leading to transversion

We analyzed energy of the eight different base pairs that results twelve base substitutions (Table 1). The optimized structures of the eight base pairs along with the A: T and G:C pairs are shown in Figure 1. In the structure, each base pair is held by hydrogen bonds (H-O and H-N). The hydrogen bond distance range was found from 1.4 to 2.2 Å. The coordinates of single bases and base pairs are given in supplementary file 1.

**Figure 1.**
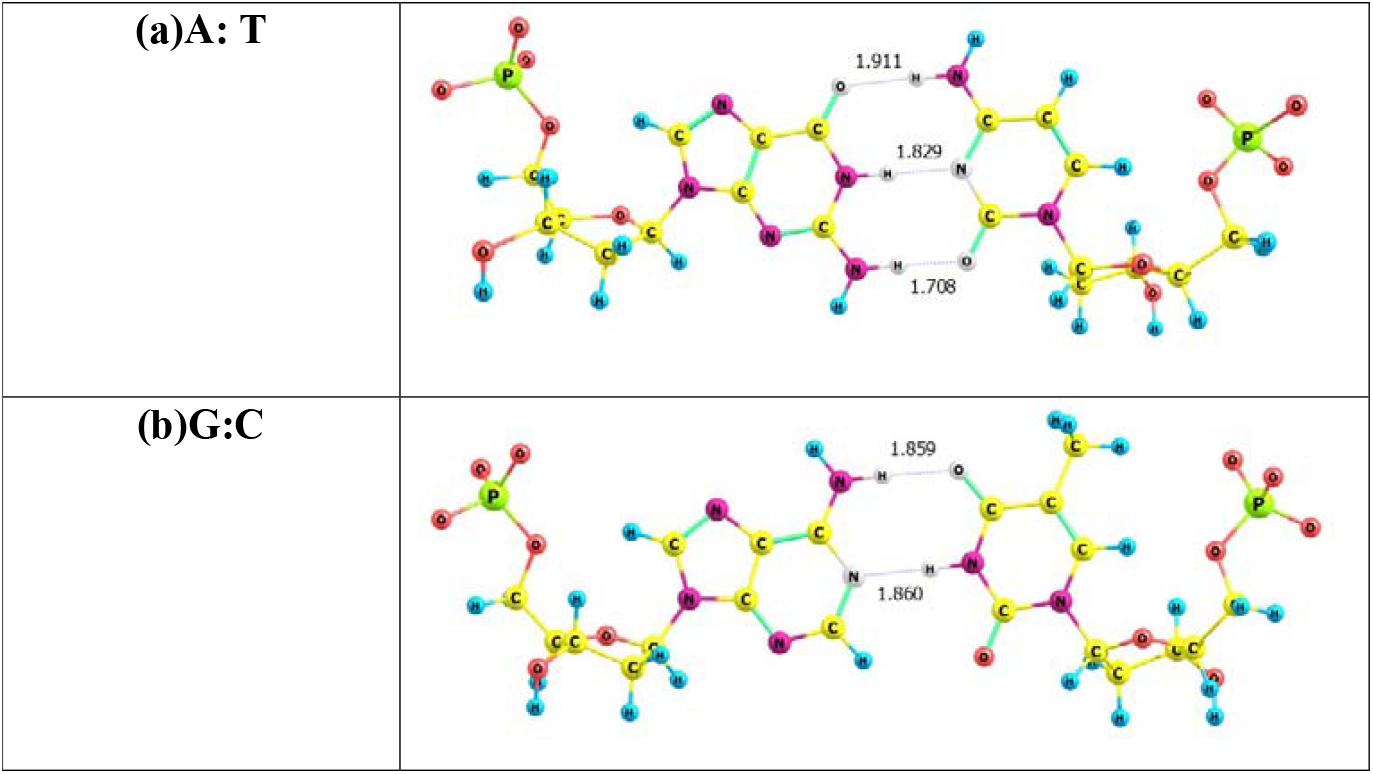
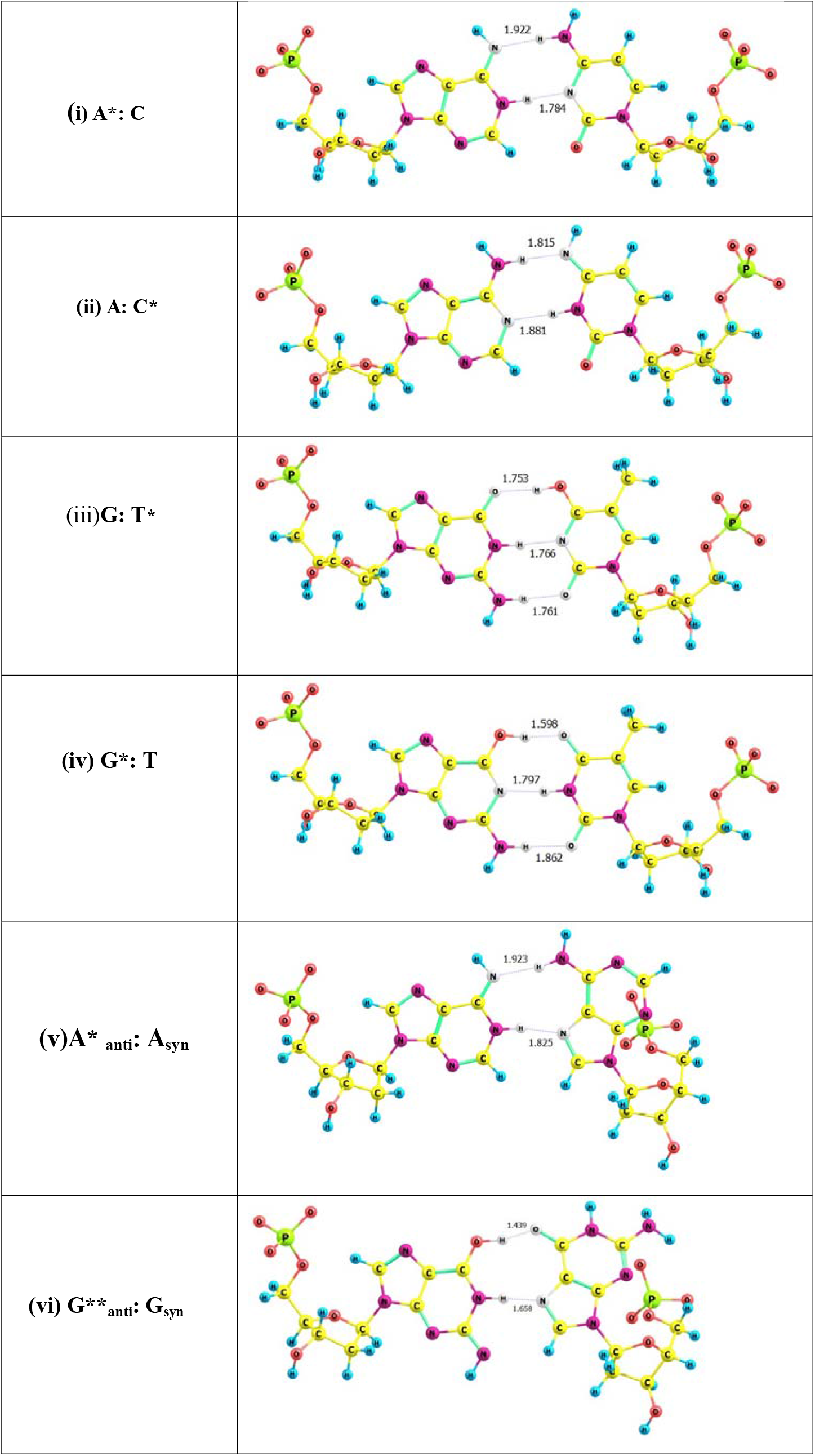
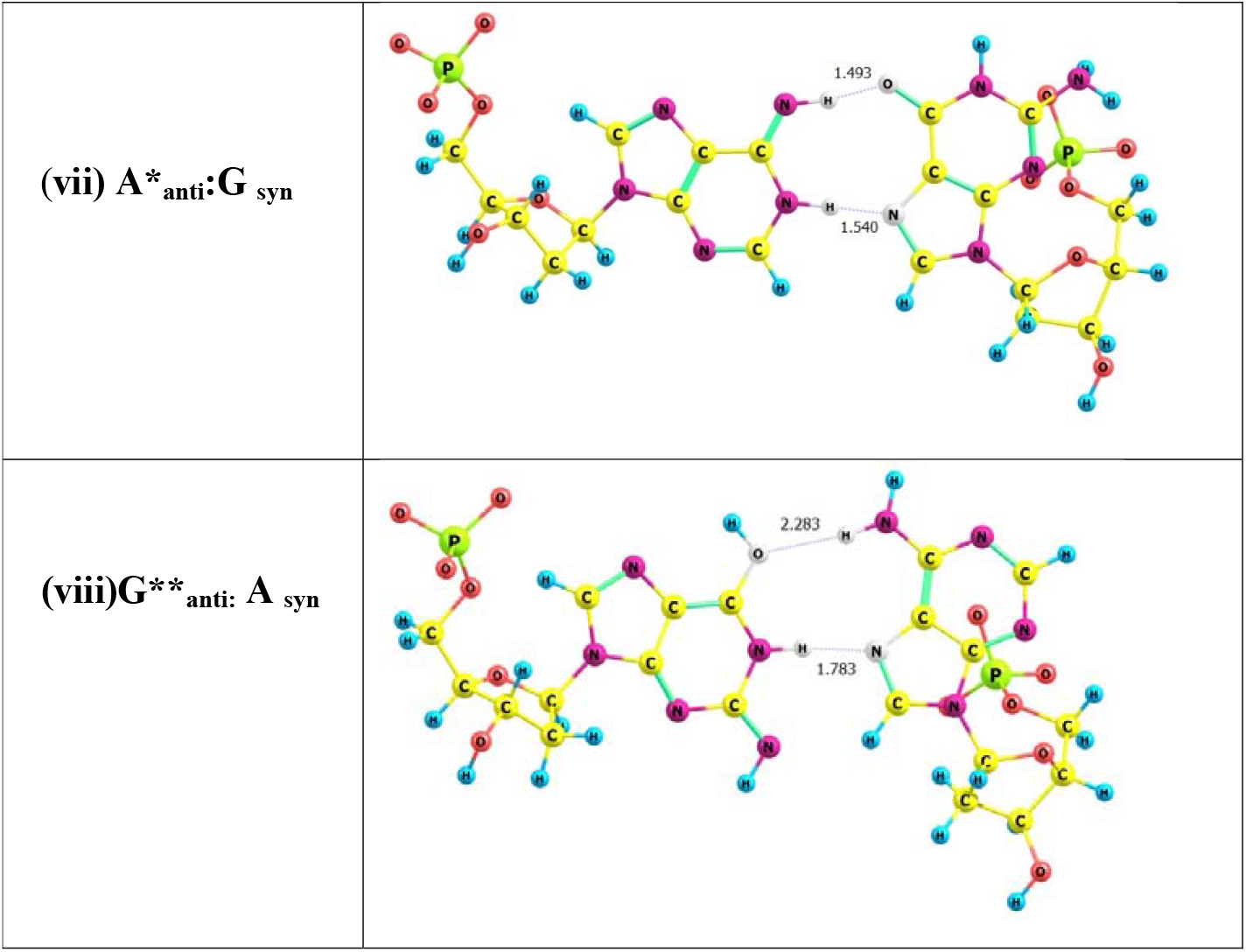
The optimized A: T and G:C pairs are showing in figure (a) and figure (b) respectively. In figures (i) to (iv) types of base pairs that can lead transition mutations are shown. Whereas, in Figure (v) to (viii) base-pairs that are responsible for transversion mutations are shown. * denotes rare tautomeric form, ** denotes the double tautomeric form.

Vibrational frequency calculations are performed to get different modes of vibrational frequency. It was found that all the single bases and base pairs have positive frequency which indicates that all molecule are in ground-state configuration. All the optimized structure corresponds to a true energy minimum on the potential energy surface (PES), meaning that the forces acting on atoms are minimized and the system is in a stable configuration. Vibrational frequencies of single bases and base pairs are given in the supplementary file 2.

Binding energy of each pair was calculated, (all calculation data is given in the supplementary file_3). Binding energy was calculated considering BSSE as well as without considering BSSE to make the observation stringent. The values of the binding energy for these ten base pairs (including A: T, G:C) are shown in Table 2.

**Table 2:**
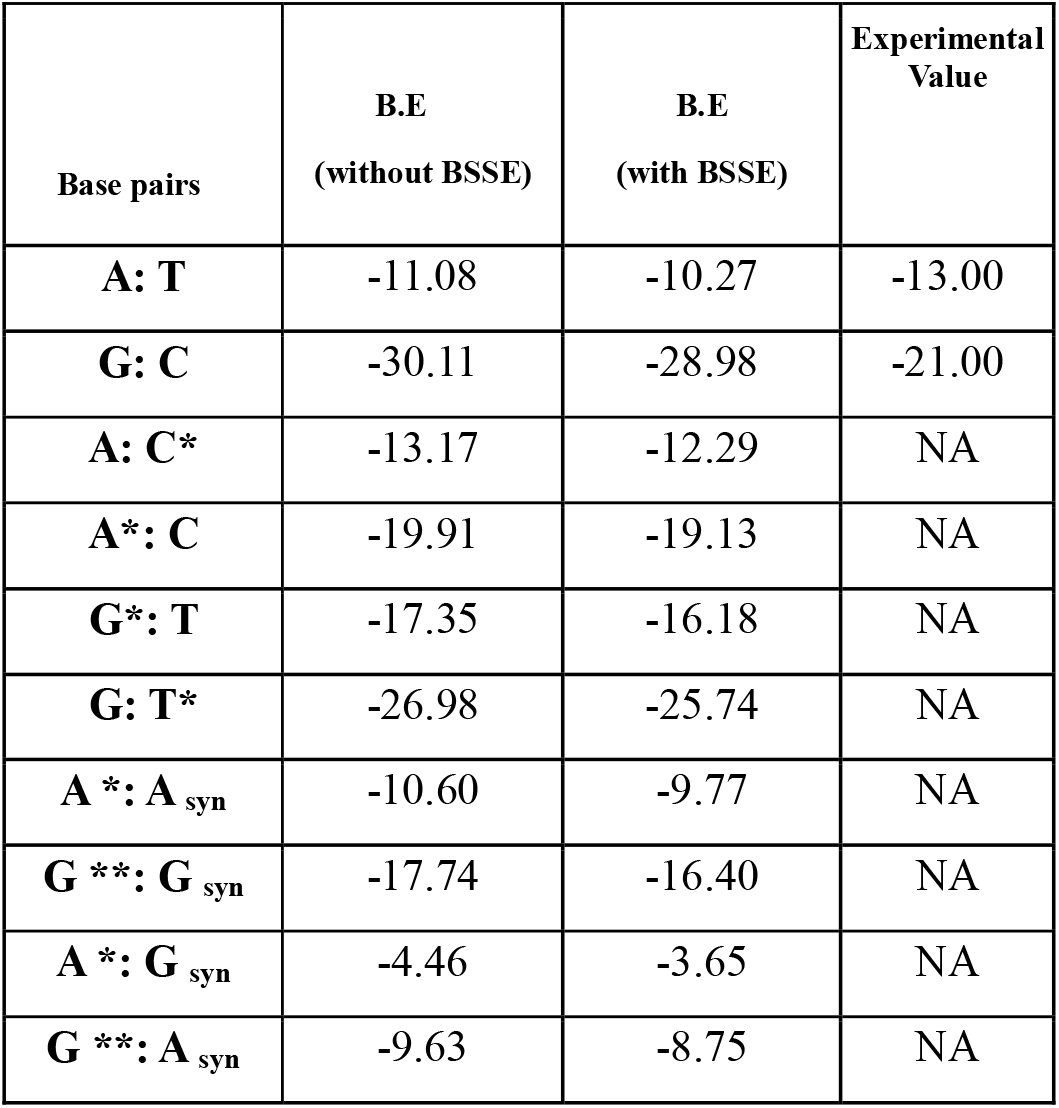
The binding Energy of DNA pairs was calculated using a 6-311++G (d, p) basis set, is shown without considering the BSSE and considering BSSE. All values are written in kcal/mole. Experimental energy value of A: T and G:C are mentioned. * denotes the rare tautomeric form., ** denotes the double tautomeric form.

The binding energies of A: T and G:C pairs were found to be -10.27 kcal/mol and - 28.98 kcal/mol, respectively. The higher stability of G:C pair than A: T is expected considering three hydrogen bonds between G and C in comparison to two hydrogen bonds between A and T. The energy calculations of these base pairs are within the range that has been reported earlier (Souri & Mohammadi, 2017; Masoodi et al., 2019, Asensio et al., 2003; Acosta-Silva et al., 2010; Thoa & Hue, 2018). The base pairs that can lead to transitions are A*(imino*): C (*amino*) (Fig. 1. (i), A(*amino*): C(*imino*) (Fig. 1. (ii)), G(*keto*): T*(enol*) (Fig. 1. (iii)), G(*enol*): T (*keto*) (Fig. 1. (iv)), whose binding energies were -19.13 kcal/mol, -12.29 kcal/mol, -25.74 kcal/mol and -16.18 kcal/mol, respectively (Table 2). A noticeable energy difference observed between the A(*imino*): C(*amino*) and A(*amino*): C(*imino*) pairs that results in G:C →A: T and A: T →G:C, respectively. Similarly, there is an energy difference between G(*keto*): T*(enol*) and G(*enol*): T (*keto*) base pairs that result in G:C →A: T and A: T →G:C transitions, respectively. So according to the binding energy, transition mutations favoring G:C → A: T is more preferred in comparison to A: T → G: C. This is interesting because by doing genome sequence analysis several researchers have observed base substitution mutation in organisms, is biased towards AT (Hershberg & Petrov, 2010),(Beura et al., 2023),(Sen et al., 2021).

Different base pairs such as A (*imino*)_anti_: A_syn_ (Fig. 1(v)), G (*enol, imino*) _anti_: G_syn_ (Fig. 1(vi)), A (*imino*)_anti_: G_syn_ (Fig. 1(vii)), and G (*enol, imino*) _anti_: A_syn_ (Fig. 1(viii)) (Topal & Fresco, 1976) can cause transversion in a genome. The binding energies of these four base pairs were estimated as -9.77 kcal/mol, -16.40 kcal/mol, -3.65 kcal/mol, and -8.75 kcal/mol (Table 2), respectively. It indicated a wide variation of their energy values. There is no available data in the literature regarding the energy values of these bases pairs that could be used as our reference to compare. However, the energy values indicate a greater magnitude of difference across different transversions occurring in organisms.

We compared the stability of base pairs leading to transition with that of base pairs leading to transversion. All four pairs resulting into transition have the energy below -10.00 Kcal/mol, whereas in the case of transversion, only one base pair has energy below -10.00 kcal/mol. This observation clearly indicates that, the base pairs that result to transition mutation are more stable than that of the base pairs that result to transversions. This observation supports the high *ti* frequency than *tv* in genomes. So far, we understand this is the first DFT comparative study of base pairs supporting the higher frequency of transition than transversion in genomes.

## 4. Discussion

Base substitution mutations in DNA are of two types such as transition and transversion. Transition results from four different base-pairs and transversion results from eight different base-pairs. However, in genomes, transition is observed to be more frequent than transversion that has been attributed to DNA geometry, RNA secondary structures, DNA repair, and codon degeneracy. However, in terms of energy of base pairs leading to these different mutations have not been reported earlier. In this study, we have calculated the stability of different possible base pairs that have the potential to cause transition as well as transversion mutations. The observations indicate that base pairs leading to *ti* are more stable than that of the base pairs leading to *tv*. So, in the present work using the DFT method we have provided another evidence in terms of binding energy in favor of the higher frequency of *ti* over *tv*. As per our knowledge, this is the first time explanation by DFT calculations regarding higher frequency of *ti* than *tv* in a genome. Though, the hypothesis by Topal and Fresco (1976) was given five decades earlier, the earlier computer programs might have been unable to run such small energy calculations, which is possible now by quantum mechanics.

Base pairs that involve tautomer such as A(*imino*):C or A:C(*imino*) and G(*enol*): T or G: T(*enol*) can result into transition mutations. Determining the stability of these base pairs experimentally, is challenging due to the short-lived nature of tautomer. However, computational studies enable accurate stability estimation by simulating experimental conditions. For transversion base pairs, both tautomer and base conformer are involved, further complicating experimental stability assessments. Quantum chemical studies, however, provide high-accuracy stability predictions.

Among the four different transitions, the C→T as well as G→A are more frequent than the other two (Kreutzer & Essigmann, 1998). Though our study on base pair stability is in support of this, there are other reasons such as frequent cytosine deamination might be attributed to the high C→T frequency (Coulondre et al., 1978). Similarly, if we observed the base pair stability in the case of *tv*, the G→T transversion was found to be less frequent. But in nature, the G→T frequency is high because of the guanine oxidation resulting in 8-oxo-guanine which causes G→T transversion(Andrés et al., 2023). Therefore, apart from the stability of base pairs, the frequency of different forms of the bases as well as the damaged bases are important attributes determining the frequency of different base substitutions.

It is pertinent to note that in this study, we analyzed energy of single base pairs only and it cannot be explained in DNA in terms of geometry, but future work could involve oligonucleotides to assess the overall stability of DNA strands with point mutations, providing insights into mutation-induced changes in DNA stability and double-helix geometry.

## Authorship contribution statement

**Nishita Deka:** Conceptualization, Methodology, Writing-Original draft preparation, Software, Visualization, Investigation. **Nand Kishor Gour**: Methodology, Data curation, Writing-Original draft preparation, Software, Validation. **Pradeep Pant:** Validation, Writing-Reviewing and Editing **Siddhartha Shankar Satapathy, Nasimul Hoda:** Reviewing and Editing, **Ramesh Chandra Deka, Suvendra Kumar Ray**: Supervision, Conceptualization, Methodology Software, Validation, Writing-Reviewing and Editing

## Acknowledgement

ND is thankful to Dr. Dikshita Dowerah for her Guidance in initial studies. ND also acknowledges DBT, GoI-funded Centre for Bioinformatics and Computational Biology, Tezpur University (BT/PR40253/BTIS/137/52/2022) for the fellowship of Scientific Administrative Assistant. ND and SKR is also thankful to the computation facility of Centre for Multidisciplinary Research (CMDR) of Tezpur University and the NNP grant awarded to SKR.

