## Supplementary File 1 for "Higher frequency of transition mutation over transversion mutation in genomes: evidence from the binding energy calculation of base pairs using DFT"

1. **Adenine**

**Co-ordinates of Single bases**

2.

X	Y	Z

15	3.364832000	-1.504110000	0.242364000

8	2.644435000	-2.464997000	1.110839000

8	4.513283000	-0.802039000	0.871107000

8	2.339822000	-0.433453000	-0.338772000

8	1.983841000	2.910740000	1.577367000

6	-0.592692000	1.805369000	-0.808551000

6	-0.232357000	2.548551000	0.482743000

6	1.200436000	2.067444000	0.763588000

6	1.753976000	1.876815000	-0.662546000

6	2.835316000	0.824476000	-0.822533000

8	0.613978000	1.539726000	-1.483741000

7	-4.499389000	-2.686357000	0.342716000

6	-4.152329000	-1.393832000	0.228760000

6	-2.839064000	-0.998139000	-0.130769000

7	-1.709790000	-1.698144000	-0.430151000

6	-0.802741000	-0.756881000	-0.682357000

7	-1.288699000	0.520968000	-0.554531000

6	-2.619499000	0.394025000	-0.201379000

7	-3.513544000	1.349696000	0.025347000

6	-4.721479000	0.836139000	0.349373000

7	-5.082674000	-0.443330000	0.463514000

1	-1.263197000	2.380983000	-1.453881000

1	-0.224851000	3.626495000	0.278162000

1	-0.937999000	2.364415000	1.293654000

1	1.185735000	1.100868000	1.273976000

1	2.168674000	2.834412000	-1.018257000

1	3.110499000	0.744717000	-1.874808000

1	3.715406000	1.129319000	-0.255910000

1	-5.442520000	-2.920536000	0.613004000

1	-3.814827000	-3.410735000	0.187406000

1	0.228693000	-0.937608000	-0.946655000

1	-5.503295000	1.565372000	0.545514000

8	3.903859000	-2.279361000	-0.922917000

1	1.992737000	3.795396000	1.185035000

**2.Thymine**

X	Y	Z

15	2.687731000	-1.283860000	0.259253000

8	1.869350000	-2.251950000	1.026482000

8	3.780559000	-0.619836000	1.015535000

8	1.743557000	-0.179701000	-0.391828000

8	1.378298000	3.288200000	1.484766000

6	-1.142155000	1.931716000	-0.828262000

6	-0.817822000	2.750503000	0.426809000

6	0.639246000	2.363093000	0.720718000

6	1.205664000	2.127262000	-0.693489000

6	2.305384000	1.080111000	-0.790411000

8	0.067609000	1.739657000	-1.511481000

8	-3.431918000	-2.896057000	0.705212000

6	-2.883192000	-1.881353000	0.309498000

7	-3.519026000	-0.629900000	0.438437000

6	-3.043917000	0.611992000	0.078616000

8	-3.662878000	1.651061000	0.226704000

7	-1.750202000	0.587866000	-0.484391000

6	-1.098970000	-0.573852000	-0.726153000

6	-1.572218000	-1.812819000	-0.344921000

6	-0.761249000	-3.040789000	-0.529470000

1	-1.868687000	2.417355000	-1.483279000

1	-0.872051000	3.813861000	0.162028000

1	-1.508741000	2.578852000	1.250490000

1	0.677988000	1.427054000	1.284956000

1	1.598877000	3.075673000	-1.089689000

1	2.682096000	1.021086000	-1.812741000

1	3.130615000	1.353815000	-0.132665000

1	-4.441472000	-0.637676000	0.859131000

1	-0.137127000	-0.464681000	-1.204214000

1	-1.352044000	-3.930416000	-0.309135000

1	-0.337042000	-3.105824000	-1.537919000

1	0.117300000	-3.015313000	0.153584000

8	3.326396000	-2.040331000	-0.868163000

1	1.326500000	4.154803000	1.05688400

**3.Guanine**

X	Y	Z

15	3.858304000	-1.291098000	0.193011000

8	3.298445000	-2.288862000	1.135110000

8	4.970270000	-0.458950000	0.720480000

8	2.693993000	-0.346375000	-0.342292000

8	2.115179000	2.975053000	1.525656000

6	-0.478481000	1.611077000	-0.706354000

6	-0.113404000	2.386520000	0.566886000

6	1.372850000	2.046272000	0.768359000

6	1.858938000	1.883102000	-0.684889000

6	3.026766000	0.937459000	-0.892707000

8	0.713639000	1.418454000	-1.430567000

7	-5.637427000	1.359742000	0.145163000

8	-4.207104000	-3.032441000	0.405572000

6	-3.848395000	-1.886960000	0.220799000

6	-2.539099000	-1.332597000	-0.037297000

7	-1.330320000	-1.940142000	-0.156990000

6	-0.478038000	-0.951673000	-0.395104000

7	-1.085731000	0.288650000	-0.430941000

6	-2.420615000	0.066663000	-0.207013000

7	-3.383026000	1.003000000	-0.174349000

6	-4.590829000	0.501804000	0.058909000

7	-4.829030000	-0.829704000	0.248235000

1	-1.212298000	2.140497000	-1.321758000

1	-0.218159000	3.460935000	0.370362000

1	-0.748396000	2.135171000	1.417734000

1	1.480431000	1.092048000	1.290381000

1	2.156430000	2.869906000	-1.077911000

1	3.236811000	0.850295000	-1.959315000

1	3.908054000	1.346057000	-0.397805000

1	-5.450157000	2.318078000	-0.107171000

1	-6.589907000	1.036243000	0.092178000

1	0.586806000	-1.058677000	-0.548222000

1	-5.763907000	-1.160448000	0.457462000

8	4.392486000	-2.047781000	-0.986617000

1	2.045049000	3.842753000	1.102798000

**4. Cytosine**

X	Y	Z

15	2.946042000	-1.340148000	0.170338000

8	2.288200000	-2.362478000	1.017494000

8	4.043979000	-0.575492000	0.816092000

8	1.854656000	-0.329041000	-0.396300000

8	1.354723000	3.036139000	1.531053000

6	-1.230083000	1.703233000	-0.731514000

6	-0.871124000	2.494928000	0.534297000

6	0.600810000	2.121604000	0.767471000

6	1.112957000	1.937297000	-0.674064000

6	2.267475000	0.967692000	-0.854242000

8	-0.029502000	1.489368000	-1.437066000

7	-3.884915000	-3.028362000	0.471554000

6	-3.213517000	-1.892142000	0.177304000

7	-3.870155000	-0.747177000	0.292115000

6	-3.250525000	0.429803000	0.006413000

8	-3.775658000	1.535134000	0.086637000

7	-1.868132000	0.387508000	-0.416780000

6	-1.215550000	-0.780167000	-0.549293000

6	-1.847172000	-1.964925000	-0.253761000

1	-1.951882000	2.226053000	-1.361609000

1	-0.950096000	3.565816000	0.309318000

1	-1.533449000	2.281959000	1.372354000

1	0.675119000	1.169175000	1.300029000

1	1.435657000	2.915171000	-1.066918000

1	2.550962000	0.925749000	-1.907062000

1	3.124047000	1.319480000	-0.278847000

1	-3.428558000	-3.924698000	0.488791000

1	-4.827905000	-2.945871000	0.819338000

1	-0.185502000	-0.729336000	-0.878674000

1	-1.314667000	-2.903271000	-0.343953000

8	3.539043000	-2.056444000	-1.007150000

1	1.254904000	3.916768000	1.142183000

**5. Adenine (imino)**

X	Y	Z

15	3.456942000	-1.461174000	0.196064000

8	2.803144000	-2.427645000	1.109852000

8	4.615842000	-0.722503000	0.760404000

8	2.378736000	-0.424128000	-0.348617000

8	2.020452000	2.918872000	1.551868000

6	-0.629545000	1.757132000	-0.725974000

6	-0.230291000	2.499335000	0.555681000

6	1.224700000	2.052417000	0.775816000

6	1.720163000	1.866077000	-0.671620000

6	2.818720000	0.838949000	-0.871087000

8	0.554348000	1.493149000	-1.439032000

7	-4.660132000	-2.721019000	0.413319000

6	-4.191467000	-1.540221000	0.232095000

6	-2.861615000	-1.057934000	-0.055259000

7	-1.703651000	-1.742663000	-0.225873000

6	-0.794206000	-0.805307000	-0.470102000

7	-1.319576000	0.469710000	-0.459969000

6	-2.657488000	0.327696000	-0.198521000

7	-3.579621000	1.313597000	-0.104717000

6	-4.783242000	0.854353000	0.152383000

1	-1.323494000	2.332094000	-1.346192000

1	-0.256596000	3.578743000	0.361190000

1	-0.894535000	2.292070000	1.395894000

1	1.255536000	1.088783000	1.290883000

1	2.094255000	2.831301000	-1.051514000

1	3.045536000	0.751705000	-1.934249000

1	3.716742000	1.173084000	-0.351030000

1	-3.904180000	-3.403229000	0.327472000

1	0.256942000	-0.976969000	-0.653972000

1	-5.606616000	1.555838000	0.247544000

8	3.959439000	-2.238662000	-0.983958000

1	-6.057857000	-0.718413000	0.516258000

1	2.011053000	3.795765000	1.142437000

7	-5.100019000	-0.452595000	0.315772000

**6.Thymine(enol)**

X	Y	Z

15	2.630877000	-1.332544000	0.263850000

8	1.774184000	-2.293027000	0.998108000

8	3.719272000	-0.702410000	1.054729000

8	1.726033000	-0.199677000	-0.393726000

8	1.406348000	3.271209000	1.510462000

6	-1.103683000	1.965435000	-0.842569000

6	-0.784038000	2.772593000	0.420747000

6	0.662221000	2.361417000	0.731794000

6	1.245771000	2.124233000	-0.675609000

6	2.323733000	1.053000000	-0.761423000

8	0.114063000	1.772897000	-1.517437000

6	-2.856728000	-1.678404000	0.292723000

7	-3.530924000	-0.574700000	0.499032000

6	-3.025257000	0.634015000	0.097825000

8	-3.597514000	1.706052000	0.225275000

7	-1.719415000	0.626035000	-0.508831000

6	-1.092817000	-0.521753000	-0.789986000

6	-1.598607000	-1.754340000	-0.394455000

1	-1.821141000	2.459550000	-1.500554000

1	-0.818295000	3.837833000	0.160144000

1	-1.489904000	2.606985000	1.232422000

1	0.678887000	1.422447000	1.292337000

1	1.669186000	3.066477000	-1.055018000

1	2.723268000	0.999630000	-1.775365000

1	3.138735000	1.300022000	-0.080760000

1	-0.143929000	-0.434835000	-1.297569000

8	3.279948000	-2.087359000	-0.858723000

8	-3.377887000	-2.828673000	0.730992000

6	-0.829187000	-3.007102000	-0.606563000

1	0.044842000	-3.023294000	0.082629000

1	-1.434790000	-3.894176000	-0.421063000

1	-0.402006000	-3.052968000	-1.614407000

1	1.361642000	4.143092000	1.092715000

1	-4.221293000	-2.608621000	1.163000000

**7.Guanine(enol)**

X	Y	Z

15	3.829206000	-1.342215000	0.112964000

8	3.252900000	-2.359281000	1.023992000

8	4.956051000	-0.546633000	0.664718000

8	2.681069000	-0.360921000	-0.390538000

8	2.225465000	2.957162000	1.557820000

6	-0.454958000	1.657941000	-0.611894000

6	-0.039835000	2.416901000	0.656905000

6	1.444842000	2.049431000	0.813400000

6	1.886119000	1.892809000	-0.654864000

6	3.035699000	0.933637000	-0.900718000

8	0.715370000	1.450729000	-1.370550000

7	-5.689015000	1.309505000	-0.095306000

8	-4.159394000	-2.978626000	0.452230000

6	-3.899562000	-1.693435000	0.227753000

6	-2.567607000	-1.252455000	0.044371000

7	-1.353064000	-1.870153000	0.028985000

6	-0.488548000	-0.889974000	-0.198403000

7	-1.083544000	0.349697000	-0.332658000

6	-2.436217000	0.142602000	-0.182704000

7	-3.409899000	1.040788000	-0.237384000

6	-4.627376000	0.475738000	-0.051101000

7	-4.906707000	-0.832022000	0.178788000

1	-1.196523000	2.205554000	-1.201388000

1	-0.130657000	3.494644000	0.471999000

1	-0.655468000	2.169655000	1.523182000

1	1.550831000	1.088648000	1.324036000

1	2.188126000	2.879667000	-1.044929000

1	3.243165000	0.876507000	-1.969860000

1	3.924468000	1.310994000	-0.394516000

1	-5.541612000	2.289598000	-0.271290000

1	-6.616732000	0.939707000	0.029497000

1	0.583057000	-1.009404000	-0.287060000

8	4.348949000	-2.070516000	-1.090733000

1	-5.123269000	-3.062895000	0.548061000

1	2.153503000	3.831360000	1.148854000

**8. Cytosine(imino)**

X	Y	Z

15	2.965217000	-1.307370000	0.163948000

8	2.330686000	-2.328718000	1.029861000

8	4.064323000	-0.525402000	0.786566000

8	1.855025000	-0.312954000	-0.395651000

8	1.334267000	3.057035000	1.513468000

6	-1.255518000	1.679761000	-0.714199000

6	-0.893954000	2.479681000	0.545778000

6	0.586677000	2.127475000	0.763966000

6	1.083926000	1.940378000	-0.682544000

6	2.247359000	0.983774000	-0.871273000

8	-0.063564000	1.469593000	-1.427281000

6	-3.182959000	-2.071844000	0.200458000

7	-3.806856000	-0.817420000	0.243954000

6	-3.246041000	0.408293000	-0.020081000

8	-3.848262000	1.465196000	0.054484000

7	-1.883495000	0.356150000	-0.391979000

6	-1.204634000	-0.821228000	-0.483566000

6	-1.794843000	-2.019364000	-0.195404000

1	-1.988246000	2.193553000	-1.340285000

1	-0.990391000	3.548725000	0.319356000

1	-1.541047000	2.260505000	1.394761000

1	0.680357000	1.179483000	1.301219000

1	1.385693000	2.919367000	-1.088407000

1	2.515853000	0.935156000	-1.927730000

1	3.108480000	1.349748000	-0.311735000

1	-0.167386000	-0.741330000	-0.783660000

8	3.548489000	-2.028184000	-1.015641000

7	-3.905912000	-3.094882000	0.510527000

1	-3.351836000	-3.949688000	0.443908000

1	-1.217227000	-2.932393000	-0.256983000

1	-4.782253000	-0.804535000	0.518339000

1	1.235800000	3.930932000	1.109231000

**9.Guanine ( imino, enol)**

X	Y	Z

15	3.634785000	-1.386051000	0.164056000

8	2.945830000	-2.678224000	0.510296000

8	4.228141000	-0.786180000	1.411019000

8	2.607063000	-0.386939000	-0.453529000

8	2.209826000	3.007558000	1.513876000

6	-0.394762000	1.681090000	-0.722398000

6	-0.024195000	2.442340000	0.557218000

6	1.455976000	2.081218000	0.765324000

6	1.946941000	1.901108000	-0.685599000

6	3.074763000	0.901252000	-0.864940000

8	0.795153000	1.477901000	-1.445704000

6	-3.801702000	-1.679983000	0.193660000

6	-2.521137000	-1.208710000	-0.061370000

7	-1.328071000	-1.862546000	-0.192604000

6	-0.459370000	-0.895023000	-0.427477000

7	-1.032176000	0.367793000	-0.453493000

6	-2.381375000	0.199384000	-0.222053000

7	-3.315152000	1.122721000	-0.164178000

6	-4.588023000	0.651063000	0.080198000

1	-1.121106000	2.223357000	-1.335352000

1	-0.113421000	3.518599000	0.364291000

1	-0.669219000	2.198743000	1.402617000

1	1.554443000	1.132070000	1.299004000

1	2.288252000	2.874626000	-1.073529000

1	3.394380000	0.851484000	-1.883879000

1	3.894828000	1.185504000	-0.241690000

1	0.601970000	-1.031516000	-0.582207000

8	4.730993000	-1.655188000	-0.832037000

1	2.130551000	3.877426000	1.096986000

7	-4.777573000	-0.760863000	0.257091000

8	-4.139287000	-2.947969000	0.379133000

7	-5.667094000	1.344411000	0.176258000

1	-5.437720000	2.329145000	0.039321000

1	-5.732967000	-1.046297000	0.441917000

1	-3.335756000	-3.493094000	0.310351000

**10.Adenine _syn_**

X	Y	Z

15	2.811696000	-1.752245000	0.042324000

8	3.979707000	-1.776041000	-0.904387000

8	1.983641000	-2.994959000	-0.149684000

8	1.933038000	-0.500832000	-0.242950000

6	2.284262000	0.436003000	-1.267441000

1	1.847993000	0.134407000	-2.194638000

1	3.350062000	0.476697000	-1.372130000

6	1.724076000	1.802113000	-0.893213000

1	2.193117000	2.563048000	-1.527756000

8	0.298752000	1.830201000	-1.137555000

6	-0.393958000	2.164460000	0.049309000

1	-0.649087000	3.233126000	0.054304000

7	-1.674818000	1.461064000	0.061342000

6	-2.892419000	2.038699000	-0.202366000

1	-2.979370000	3.107916000	-0.344383000

7	-3.903891000	1.189639000	-0.261326000

6	-3.313550000	-0.026601000	-0.044224000

6	-3.828813000	-1.347552000	0.001656000

7	-2.984463000	-2.377014000	0.212943000

6	-1.688023000	-2.103602000	0.387007000

7	-1.079987000	-0.899017000	0.388439000

6	-1.924801000	0.095540000	0.156915000

6	1.912520000	2.192148000	0.585563000

1	2.736229000	1.637379000	1.049183000

6	0.555837000	1.825533000	1.201901000

1	0.518317000	0.755558000	1.411623000

8	2.149651000	3.599719000	0.611407000

1	2.337048000	3.851675000	1.525274000

1	0.334110000	2.392889000	2.109978000

8	3.313384000	-1.698381000	1.459740000

7	-5.138248000	-1.607180000	-0.163861000

1	-5.451862000	-2.565627000	-0.142709000

1	-5.787193000	-0.858320000	-0.348551000

1	-1.031132000	-2.955608000	0.538958000

**11.Guanine _syn_**

X	Y	Z

15	-2.775966000	-1.660204000	-0.144148000

8	-3.884957000	-1.747383000	0.869778000

8	-1.891614000	-2.875205000	-0.034609000

8	-1.932618000	-0.379709000	0.129203000

6	-2.266174000	0.469809000	1.231889000

1	-1.761182000	0.118524000	2.106690000

1	-3.324507000	0.442204000	1.392830000

6	-1.765394000	1.877690000	0.935993000

1	-2.255738000	2.583788000	1.614724000

8	-0.336333000	1.952517000	1.182270000

6	0.341533000	2.340206000	0.010155000

1	0.588604000	3.410277000	0.040356000

7	1.640901000	1.654798000	-0.036930000

6	2.865072000	2.268333000	-0.002870000

1	2.950310000	3.346501000	0.023128000

7	3.897983000	1.432823000	-0.003573000

6	3.328405000	0.205142000	-0.017299000

6	3.938497000	-1.113219000	-0.019074000

7	2.928301000	-2.124417000	-0.023789000

6	1.567084000	-1.922524000	-0.023574000

7	1.028107000	-0.690074000	-0.047634000

6	1.910319000	0.294064000	-0.033587000

6	-1.967258000	2.342217000	-0.517871000

1	-2.781589000	1.796282000	-1.006478000

6	-0.604526000	2.032526000	-1.155223000

1	-0.550454000	0.974507000	-1.415397000

8	-2.223997000	3.745001000	-0.471789000

1	-2.453010000	4.035145000	-1.364533000

1	-0.397743000	2.644763000	-2.037044000

8	-3.367652000	-1.597662000	-1.527020000

8	5.113357000	-1.422217000	-0.014043000

7	0.739036000	-2.961420000	0.008917000

1	3.301970000	-3.066600000	-0.007178000

1	-0.311284000	-2.827578000	0.009066000

1	1.098982000	-3.904921000	0.00557100

**Co-ordinates of Base Pairs**

**A: T pair**

X	Y	Z

15	-9.424452000	1.781554000	-0.017808000

8	-8.772912000	2.803771000	0.834889000

8	-10.691061000	1.212814000	0.510772000

8	-8.397928000	0.603268000	-0.322048000

8	-8.351535000	-2.401595000	2.028323000

6	-5.559768000	-1.911540000	-0.329180000

6	-6.053314000	-2.369281000	1.051780000

6	-7.460251000	-1.753490000	1.150290000

6	-7.909410000	-1.754824000	-0.323915000

6	-8.899406000	-0.678598000	-0.731028000

8	-6.701329000	-1.606274000	-1.097959000

7	-1.020907000	2.117282000	-0.242305000

6	-1.534969000	0.886965000	-0.220408000

6	-2.933180000	0.628164000	-0.259886000

7	-4.018547000	1.445369000	-0.339252000

6	-5.055281000	0.609754000	-0.357971000

7	-4.693053000	-0.712974000	-0.284159000

6	-3.312802000	-0.731642000	-0.226997000

7	-2.502644000	-1.783444000	-0.164154000

6	-1.205703000	-1.417227000	-0.137061000

7	-0.696145000	-0.181978000	-0.159863000

1	-4.953267000	-2.673711000	-0.827552000

1	-6.132603000	-3.463335000	1.053915000

1	-5.382537000	-2.080267000	1.862385000

1	-7.403756000	-0.721077000	1.504595000

1	-8.364432000	-2.730738000	-0.562344000

1	-9.019944000	-0.710994000	-1.814445000

1	-9.867180000	-0.873135000	-0.268301000

1	-0.002963000	2.261814000	-0.210861000

1	-1.650112000	2.906157000	-0.288120000

1	-6.093461000	0.905235000	-0.436736000

1	-0.474229000	-2.220005000	-0.089553000

8	-9.746286000	2.437939000	-1.327398000

15	9.121701000	1.578233000	0.178407000

8	8.492369000	2.625844000	-0.659471000

8	10.375636000	0.989069000	-0.357963000

8	8.070035000	0.418698000	0.468263000

8	8.043947000	-2.660303000	-1.869691000

6	5.172520000	-2.002212000	0.350138000

6	5.708684000	-2.522936000	-0.994626000

6	7.138373000	-1.954640000	-1.053513000

6	7.524048000	-1.921574000	0.437785000

6	8.543722000	-0.879222000	0.858994000

8	6.290651000	-1.688791000	1.147822000

8	1.849576000	2.396841000	-0.125750000

6	2.641031000	1.451886000	-0.027747000

7	2.181695000	0.141704000	-0.027784000

6	2.935210000	-1.004851000	0.070328000

8	2.456220000	-2.129594000	0.036668000

7	4.318008000	-0.784271000	0.211441000

6	4.847596000	0.473949000	0.228445000

6	4.088063000	1.608194000	0.098622000

6	4.679908000	2.981471000	0.106109000

1	4.531921000	-2.729074000	0.851402000

1	5.751612000	-3.618286000	-0.956915000

1	5.081786000	-2.243653000	-1.842582000

1	7.132460000	-0.933997000	-1.445652000

1	7.920806000	-2.909113000	0.727174000

1	8.664257000	-0.928832000	1.942306000

1	9.507075000	-1.088652000	0.393548000

1	1.153757000	-0.002153000	-0.100161000

1	5.922147000	0.525140000	0.366809000

1	4.212121000	3.598840000	-0.666331000

1	4.487626000	3.485622000	1.062230000

1	5.760585000	2.953501000	-0.055240000

8	9.458526000	2.209729000	1.497108000

1	-8.414791000	-3.332968000	1.772965000

1	8.057832000	-3.586360000	-1.58846300

**G: C Pair**

X	Y	Z

15	9.310318000	1.831559000	0.082393000

8	8.662131000	2.867508000	-0.756165000

8	10.572471000	1.262452000	-0.456344000

8	8.278265000	0.655228000	0.375049000

8	8.222278000	-2.343680000	-2.000407000

6	5.414367000	-1.830299000	0.331598000

6	5.917093000	-2.287399000	-1.046387000

6	7.333166000	-1.686001000	-1.128830000

6	7.768103000	-1.694147000	0.348969000

6	8.773459000	-0.634127000	0.767215000

8	6.548196000	-1.523809000	1.107679000

7	0.146206000	-2.307777000	0.046434000

8	0.909409000	2.229952000	0.142378000

6	1.402012000	1.107269000	0.150972000

6	2.812422000	0.766086000	0.219184000

7	3.904804000	1.546929000	0.305358000

6	4.932545000	0.688889000	0.344855000

7	4.547853000	-0.624754000	0.276369000

6	3.166442000	-0.617325000	0.203560000

7	2.362014000	-1.665358000	0.142429000

6	1.044508000	-1.330153000	0.094760000

7	0.590403000	-0.041730000	0.091343000

1	4.801023000	-2.588784000	0.827821000

1	5.985850000	-3.382039000	-1.052418000

1	5.259317000	-1.988170000	-1.864122000

1	7.292128000	-0.652303000	-1.481653000

1	8.198943000	-2.678916000	0.594973000

1	8.897126000	-0.680399000	1.849825000

1	9.739358000	-0.828292000	0.299712000

1	0.504735000	-3.251405000	0.049100000

1	-0.884340000	-2.132198000	0.015181000

1	5.974995000	0.969023000	0.435218000

1	-0.447715000	0.125620000	0.036953000

8	9.639057000	2.470317000	1.398974000

15	-9.237107000	1.730390000	-0.067446000

8	-8.599968000	2.764391000	0.781362000

8	-10.492831000	1.141537000	0.465056000

8	-8.192529000	0.568814000	-0.374020000

8	-8.027093000	-2.503542000	1.936386000

6	-5.261764000	-1.794557000	-0.404578000

6	-5.734455000	-2.318620000	0.960462000

6	-7.172461000	-1.779937000	1.079385000

6	-7.625621000	-1.756855000	-0.393061000

6	-8.673818000	-0.721650000	-0.779204000

8	-6.423632000	-1.508607000	-1.156105000

7	-1.974944000	2.682530000	-0.111762000

6	-2.795830000	1.625054000	-0.176417000

7	-2.251276000	0.397860000	-0.095274000

6	-3.034320000	-0.711754000	-0.144808000

8	-2.570938000	-1.867493000	-0.049668000

7	-4.422809000	-0.557546000	-0.308859000

6	-4.976032000	0.685159000	-0.404826000

6	-4.204898000	1.802268000	-0.331918000

1	-4.633928000	-2.513482000	-0.932615000

1	-5.759338000	-3.415148000	0.930819000

1	-5.078718000	-2.022882000	1.780367000

1	-7.170297000	-0.758311000	1.468328000

1	-8.019168000	-2.750623000	-0.665068000

1	-8.797963000	-0.757667000	-1.862687000

1	-9.637497000	-0.931568000	-0.313867000

1	-2.350464000	3.615031000	-0.168900000

1	-0.966606000	2.563194000	-0.007723000

1	-6.049292000	0.720702000	-0.548297000

1	-4.652241000	2.785367000	-0.401364000

8	-9.573893000	2.379596000	-1.377538000

1	8.294673000	-3.270183000	-1.730080000

1	-8.031387000	-3.429590000	1.655336000

**A*:C Pair**

X	Y	Z

15	-9.298046000	1.726349000	-0.252765000

8	-8.618321000	2.876886000	0.388113000

8	-10.534646000	1.251405000	0.419803000

8	-8.275679000	0.515453000	-0.403920000

8	-8.149281000	-2.136386000	2.380442000

X	Y	Z

6	-5.394933000	-1.924653000	-0.064343000

6	-5.866499000	-2.206256000	1.371248000

6	-7.281705000	-1.598973000	1.409867000

6	-7.750259000	-1.801378000	-0.043028000

6	-8.782729000	-0.817340000	-0.571562000

8	-6.550627000	-1.715330000	-0.845501000

7	-0.886231000	2.210428000	-0.350885000

6	-1.417257000	1.031866000	-0.283226000

6	-2.810940000	0.668456000	-0.283083000

7	-3.914723000	1.445449000	-0.372357000

6	-4.930724000	0.586146000	-0.308354000

7	-4.528901000	-0.722865000	-0.174530000

6	-3.150448000	-0.700068000	-0.164175000

7	-2.300712000	-1.732630000	-0.068968000

6	-1.023277000	-1.355829000	-0.095108000

1	-4.792193000	-2.739017000	-0.477152000

1	-5.932597000	-3.291560000	1.515413000

1	-5.190669000	-1.808530000	2.130128000

1	-7.235386000	-0.527899000	1.624077000

1	-8.173311000	-2.814623000	-0.147120000

1	-8.951983000	-1.029315000	-1.627881000

1	-9.727664000	-0.933654000	-0.039420000

1	-1.625033000	2.912862000	-0.418333000

1	-5.978837000	0.850498000	-0.373439000

1	-0.242066000	-2.108976000	-0.038535000

8	-9.683879000	2.155773000	-1.637068000

15	9.204682000	1.636228000	0.438545000

8	8.637313000	2.765494000	-0.335012000

8	10.513279000	1.119443000	-0.038525000

8	8.151537000	0.443988000	0.499289000

8	8.133533000	-2.353211000	-2.160115000

6	5.224259000	-1.867562000	0.066497000

6	5.784012000	-2.237314000	-1.318749000

6	7.233288000	-1.713097000	-1.283753000

6	7.589419000	-1.860513000	0.208025000

6	8.609389000	-0.886001000	0.788431000

8	6.342359000	-1.679090000	0.914914000

6	2.798354000	1.586422000	0.006596000

7	2.226925000	0.379315000	-0.095249000

6	2.977596000	-0.764307000	-0.079425000

8	2.486360000	-1.888581000	-0.183322000

7	4.394351000	-0.627364000	0.061501000

6	4.966547000	0.595547000	0.200096000

6	4.214196000	1.732096000	0.170072000

1	4.561986000	-2.636229000	0.466564000

1	5.796172000	-3.330008000	-1.416031000

1	5.187676000	-1.837446000	-2.140190000

1	7.268063000	-0.655529000	-1.558167000

1	7.958393000	-2.883657000	0.390980000

1	8.630339000	-1.054068000	1.866165000

1	9.615800000	-1.034906000	0.395481000

1	6.039895000	0.613413000	0.349702000

8	9.406318000	2.120280000	1.844386000

7	2.008185000	2.668694000	-0.041735000

1	2.414692000	3.586964000	0.034033000

1	0.992469000	2.573486000	-0.169177000

1	0.469995000	0.078767000	-0.179318000

1	4.681181000	2.703059000	0.274264000

1	-8.213498000	-3.092436000	2.244156000

1	8.107299000	-3.304919000	-1.986216000

7	-0.578340000	-0.090656000	-0.187878000

**A: C* pair**

X	Y	Z

15	9.309939000	1.774157000	0.082371000

8	8.663815000	2.812573000	-0.754732000

8	10.571345000	1.203822000	-0.456829000

8	8.275766000	0.599068000	0.372707000

8	8.226082000	-2.396130000	-2.008106000

6	5.422283000	-1.895583000	0.332774000

6	5.922680000	-2.357257000	-1.044248000

6	7.332926000	-1.747448000	-1.132613000

6	7.772820000	-1.757490000	0.344108000

6	8.768967000	-0.691825000	0.762912000

8	6.560656000	-1.604621000	1.111398000

7	0.945390000	2.200011000	0.138432000

6	1.438052000	0.964122000	0.151280000

6	2.832395000	0.681935000	0.215703000

7	3.927914000	1.482652000	0.293199000

6	4.952505000	0.631071000	0.335227000

7	4.571142000	-0.686050000	0.277672000

6	3.190445000	-0.685050000	0.208135000

7	2.363943000	-1.722642000	0.154315000

6	1.070423000	-1.335045000	0.108675000

1	4.801609000	-2.649623000	0.825847000

1	5.997631000	-3.451635000	-1.044296000

1	5.258798000	-2.066498000	-1.859914000

1	7.283464000	-0.713283000	-1.483258000

1	8.219419000	-2.737799000	0.580653000

1	8.891698000	-0.739949000	1.845504000

1	9.734404000	-0.886767000	0.295384000

1	1.597887000	2.971475000	0.172935000

1	5.994114000	0.912825000	0.421015000

1	0.327490000	-2.127766000	0.071373000

8	9.639240000	2.410185000	1.400124000

15	-9.237569000	1.704336000	-0.074776000

8	-8.599026000	2.735862000	0.775989000

8	-10.494484000	1.116708000	0.456275000

8	-8.194785000	0.541514000	-0.382900000

8	-8.083399000	-2.532002000	1.930786000

6	-5.298665000	-1.874402000	-0.396276000

6	-5.783664000	-2.391519000	0.968011000

6	-7.211693000	-1.826430000	1.077869000

6	-7.654834000	-1.798645000	-0.397919000

6	-8.678406000	-0.747542000	-0.790364000

8	-6.446279000	-1.578012000	-1.156850000

6	-2.761631000	1.589146000	-0.169328000

7	-2.306327000	0.280668000	-0.077760000

6	-3.068216000	-0.860937000	-0.118401000

8	-2.600114000	-1.985473000	-0.011430000

7	-4.451244000	-0.647884000	-0.296101000

6	-4.971843000	0.609359000	-0.410551000

6	-4.188715000	1.722259000	-0.341381000

1	-4.670713000	-2.601287000	-0.914579000

1	-5.824777000	-3.487191000	0.935123000

1	-5.126351000	-2.108220000	1.790951000

1	-7.193188000	-0.804062000	1.464728000

1	-8.067582000	-2.784291000	-0.669513000

1	-8.801738000	-0.781410000	-1.873972000

1	-9.642291000	-0.956542000	-0.325673000

1	-6.043451000	0.663587000	-0.564888000

8	-9.572759000	2.356267000	-1.383917000

7	0.581111000	-0.093994000	0.101992000

7	-1.882407000	2.543836000	-0.096445000

1	-2.334010000	3.453144000	-0.178884000

1	-4.635130000	2.704694000	-0.423177000

1	-0.084080000	2.367825000	0.069423000

1	-1.284791000	0.127570000	0.020067000

1	8.284936000	-3.328446000	-1.755126000

1	-8.099010000	-3.460322000	1.657419000

**G*: T pair**

X	Y	Z

15	9.409266000	1.920704000	0.042911000

8	8.759230000	2.941764000	-0.812303000

8	10.676369000	1.350647000	-0.483029000

8	8.381966000	0.743314000	0.347728000

8	8.394063000	-2.253020000	-2.005137000

6	5.570401000	-1.836186000	0.323943000

6	6.086865000	-2.280370000	-1.052983000

6	7.479355000	-1.629238000	-1.135314000

6	7.910495000	-1.623576000	0.343536000

6	8.882177000	-0.538091000	0.759562000

8	6.690041000	-1.492856000	1.102126000

7	0.378927000	-2.561453000	0.004764000

8	0.921803000	2.003902000	0.200938000

6	1.431913000	0.808955000	0.175795000

6	2.852883000	0.623719000	0.231453000

7	3.902145000	1.468863000	0.331165000

6	4.975662000	0.669990000	0.357513000

7	4.667871000	-0.662821000	0.267353000

6	3.291714000	-0.734086000	0.192409000

7	2.549341000	-1.818598000	0.113260000

6	1.215537000	-1.520550000	0.075215000

7	0.660694000	-0.280157000	0.100562000

1	4.980087000	-2.613666000	0.818839000

1	6.193783000	-3.371966000	-1.055746000

1	5.417898000	-2.006381000	-1.870401000

1	7.401478000	-0.597205000	-1.486938000

1	8.374749000	-2.593554000	0.588647000

1	9.001573000	-0.568784000	1.843103000

1	9.850692000	-0.733676000	0.298341000

1	0.784256000	-3.485193000	-0.015260000

1	-0.642609000	-2.443615000	-0.029251000

1	6.000249000	1.008008000	0.452422000

8	9.729598000	2.579127000	1.351872000

15	-9.136682000	1.723779000	-0.177617000

8	-8.508084000	2.772528000	0.659402000

8	-10.391544000	1.135941000	0.358074000

8	-8.085047000	0.563377000	-0.464140000

8	-8.116481000	-2.486145000	1.888334000

6	-5.214117000	-1.947494000	-0.324775000

6	-5.772232000	-2.429299000	1.025295000

6	-7.183052000	-1.816811000	1.071731000

6	-7.556743000	-1.787528000	-0.421943000

6	-8.558598000	-0.734964000	-0.853253000

8	-6.316829000	-1.585533000	-1.125201000

6	-2.540724000	1.392583000	0.007444000

6	-2.948666000	-1.041750000	-0.105234000

8	-2.486652000	-2.191083000	-0.093298000

7	-4.299819000	-0.773209000	-0.205083000

6	-4.773069000	0.525299000	-0.195370000

6	-3.970214000	1.619354000	-0.079517000

1	-4.614506000	-2.711555000	-0.821575000

1	-5.852210000	-3.523277000	1.007761000

1	-5.139160000	-2.155275000	1.870608000

1	-7.148411000	-0.792689000	1.452976000

1	-7.972965000	-2.769937000	-0.704304000

1	-8.677831000	-0.786346000	-1.936768000

1	-9.522787000	-0.943367000	-0.388988000

1	-5.847292000	0.617761000	-0.309493000

8	-9.471527000	2.353225000	-1.497775000

8	-1.675323000	2.283421000	0.095744000

6	-4.497082000	3.025639000	-0.057594000

1	-5.587431000	3.037349000	0.006455000

1	-4.090437000	3.572272000	0.798593000

1	-4.191672000	3.573599000	-0.955598000

1	-1.127392000	-0.109497000	0.039706000

1	-0.098474000	2.030677000	0.158232000

1	8.496259000	-3.177725000	-1.738306000

1	-8.162154000	-3.411927000	1.609892000

7	-2.148036000	0.065416000	-0.015810000

**G: T ***

X	Y	Z

15	9.415019000	1.894210000	0.054547000

8	8.760474000	2.920442000	-0.791001000

8	10.682934000	1.332402000	-0.478310000

8	8.391719000	0.711182000	0.351152000

8	8.379748000	-2.301172000	-1.995981000

6	5.554996000	-1.836566000	0.326574000

6	6.070364000	-2.288050000	-1.049413000

6	7.473324000	-1.657277000	-1.132245000

6	7.902503000	-1.645572000	0.347102000

6	8.896725000	-0.572082000	0.751292000

8	6.678527000	-1.478345000	1.097014000

7	0.316129000	-2.540427000	0.048139000

8	0.864067000	2.013836000	0.164748000

6	1.424990000	0.917915000	0.162014000

6	2.849282000	0.648273000	0.215268000

7	3.909944000	1.475780000	0.291002000

6	4.972457000	0.663326000	0.323675000

7	4.642853000	-0.666710000	0.266178000

6	3.263452000	-0.718888000	0.201086000

7	2.508737000	-1.806126000	0.144018000

6	1.181267000	-1.530683000	0.099943000

7	0.673475000	-0.263823000	0.104273000

1	4.972699000	-2.614340000	0.830508000

1	6.162023000	-3.380978000	-1.052440000

1	5.407546000	-2.005221000	-1.868879000

1	7.411164000	-0.626788000	-1.491414000

1	8.343537000	-2.622968000	0.604917000

1	9.021453000	-0.581593000	1.840116000

1	9.864844000	-0.760939000	0.286931000

1	0.699665000	-3.473784000	0.042777000

1	-0.711622000	-2.386741000	0.009452000

1	6.002964000	0.987002000	0.403193000

1	-0.370849000	-0.124384000	0.061870000

8	9.735596000	2.542574000	1.368427000

15	-9.130602000	1.643233000	-0.136634000

8	-8.503732000	2.686816000	0.708087000

8	-10.382783000	1.047148000	0.396169000

8	-8.075984000	0.488469000	-0.434697000

8	-7.990467000	-2.602911000	1.877645000

6	-5.145728000	-1.905594000	-0.369526000

6	-5.665832000	-2.435948000	0.976947000

6	-7.100578000	-1.881314000	1.056379000

6	-7.503637000	-1.839521000	-0.429516000

6	-8.546396000	-0.808027000	-0.834029000

8	-6.278700000	-1.577389000	-1.146646000

6	-2.626751000	1.438108000	-0.023106000

7	-2.101903000	0.215430000	-0.022670000

6	-2.897280000	-0.890961000	-0.104137000

8	-2.437964000	-2.049556000	-0.066450000

7	-4.276214000	-0.693963000	-0.233439000

6	-4.809249000	0.564830000	-0.235348000

6	-4.032811000	1.683039000	-0.112437000

1	-4.520722000	-2.631701000	-0.890891000

1	-5.702755000	-3.531690000	0.937729000

1	-5.032446000	-2.154559000	1.819399000

1	-7.099778000	-0.863626000	1.455924000

1	-7.885402000	-2.831595000	-0.725154000

1	-8.667183000	-0.850633000	-1.917605000

1	-9.509011000	-1.023145000	-0.369664000

1	-5.885060000	0.620184000	-0.357781000

8	-9.469571000	2.282738000	-1.450923000

8	-1.835101000	2.489482000	0.057984000

6	-4.596980000	3.076443000	-0.093274000

1	-5.687211000	3.053487000	-0.027893000

1	-4.211566000	3.637814000	0.762933000

1	-4.311595000	3.632374000	-0.992837000

1	8.470358000	-3.224495000	-1.720288000

1	-8.000952000	-3.525178000	1.584461000

1	-0.872311000	2.246285000	0.102196000

**A*: A _Syn_ pair**

X	Y	Z

15	8.983136000	1.134184000	-0.538216000

8	8.387047000	2.457995000	-0.942800000

8	9.399682000	0.376222000	-1.770616000

8	7.927184000	0.302732000	0.252015000

8	7.112912000	-3.163275000	-1.327582000

6	4.823397000	-1.701007000	1.159697000

6	5.028035000	-2.576884000	-0.085460000

6	6.447549000	-2.202553000	-0.540951000

6	7.128563000	-1.912644000	0.808255000

6	8.312029000	-0.971742000	0.775300000

8	6.099041000	-1.379088000	1.660505000

7	0.601642000	2.544086000	-0.051447000

6	1.106873000	1.388784000	0.236366000

6	2.477053000	0.991343000	0.442236000

7	3.631740000	1.706053000	0.366721000

6	4.590576000	0.821765000	0.618226000

7	4.101140000	-0.444901000	0.859029000

6	2.739377000	-0.355048000	0.752122000

7	1.851812000	-1.361194000	0.902386000

6	0.615724000	-0.959259000	0.696689000

1	4.226798000	-2.199656000	1.929223000

1	5.007936000	-3.631858000	0.214887000

1	4.257664000	-2.428450000	-0.843199000

1	6.436565000	-1.289753000	-1.143081000

1	7.487463000	-2.865420000	1.234434000

1	8.706598000	-0.840890000	1.761071000

1	9.058608000	-1.400390000	0.141347000

1	1.350602000	3.232876000	-0.132707000

1	5.651560000	1.030456000	0.639228000

1	-0.189411000	-1.682885000	0.782738000

8	10.185941000	1.372887000	0.335708000

1	7.121921000	-4.003226000	-0.846760000

7	0.234375000	0.295973000	0.379190000

15	-6.280823000	0.511278000	2.043806000

8	-7.667823000	0.685913000	2.601302000

8	-5.589672000	1.848471000	1.975896000

8	-6.370651000	-0.097482000	0.613452000

6	-7.633825000	-0.400089000	0.000770000

1	-7.947969000	0.421391000	-0.608573000

1	-8.374174000	-0.582925000	0.754157000

6	-7.448095000	-1.586958000	-0.929131000

1	-8.404574000	-1.828760000	-1.400865000

8	-6.537665000	-1.245701000	-2.002868000

6	-5.250296000	-1.663675000	-1.704903000

1	-4.821918000	-2.120240000	-2.603527000

7	-4.340547000	-0.505251000	-1.380021000

6	-3.080705000	-0.691449000	-0.913680000

1	-2.657642000	-1.681382000	-0.811989000

7	-2.436909000	0.427766000	-0.601534000

6	-3.336261000	1.416550000	-0.877046000

6	-3.258232000	2.839120000	-0.795094000

7	-2.160948000	3.497289000	-0.417500000

1	-2.234867000	4.505974000	-0.399995000

1	-1.239923000	3.060936000	-0.246415000

7	-4.351850000	3.559258000	-1.139364000

6	-5.427951000	2.908804000	-1.565748000

1	-6.281035000	3.528841000	-1.828591000

7	-5.607892000	1.578122000	-1.741236000

6	-4.548813000	0.874056000	-1.373161000

6	-6.844566000	-2.845358000	-0.295907000

1	-7.093732000	-2.916803000	0.770347000

6	-5.330719000	-2.642747000	-0.502995000

1	-4.890867000	-2.193398000	0.388229000

8	-7.351088000	-3.958793000	-1.021991000

1	-7.060236000	-4.765640000	-0.577450000

1	-4.831929000	-3.591532000	-0.718872000

1	-0.764132000	0.451944000	0.128739000

8	-5.640436000	-0.244582000	2.772815000

**A*: G _syn_ pair**

X	Y	Z

15	9.029209000	1.585582000	-0.379561000

8	8.353729000	2.742000000	-1.062010000

8	9.547031000	0.620806000	-1.411063000

8	8.008867000	0.856995000	0.538130000

8	7.919127000	-2.351061000	-0.928870000

6	5.129100000	-1.836895000	1.381484000

6	5.853395000	-2.758506000	0.390188000

6	6.929434000	-1.802681000	-0.068418000

6	7.420922000	-1.407989000	1.330555000

6	8.460998000	-0.296276000	1.235236000

8	6.195627000	-1.154504000	2.113294000

7	0.883331000	1.638952000	-1.015583000

6	1.292794000	0.474284000	-0.595604000

6	2.624987000	0.308655000	-0.159738000

7	3.689850000	1.238564000	-0.125201000

6	4.674814000	0.530612000	0.401789000

7	4.326796000	-0.776181000	0.735088000

6	3.037598000	-0.919370000	0.353621000

7	2.257890000	-2.023664000	0.427153000

6	1.032686000	-1.876189000	-0.038308000

1	4.439465000	-2.390965000	2.016618000

1	6.305922000	-3.595503000	0.918225000

1	5.217469000	-3.162812000	-0.395559000

1	6.507514000	-0.957518000	-0.587238000

1	7.899147000	-2.276876000	1.783630000

1	8.871858000	-0.004293000	2.178533000

1	9.217047000	-0.779470000	0.649180000

1	5.655469000	0.944697000	0.588707000

1	0.369149000	-2.738912000	-0.023050000

8	10.177945000	2.091986000	0.452063000

1	8.320484000	-3.117038000	-0.499248000

7	0.520563000	-0.692916000	-0.499428000

15	-6.473759000	1.500404000	2.183184000

8	-7.903812000	1.774246000	2.566150000

8	-5.769476000	2.797088000	1.882041000

8	-6.438455000	0.583069000	0.926685000

6	-7.655474000	0.127027000	0.326268000

1	-7.953536000	0.798277000	-0.449956000

1	-8.430045000	0.083765000	1.068388000

6	-7.596189000	-1.336239000	-0.452080000

1	-8.609393000	-1.344399000	-0.863350000

8	-6.524442000	-1.219970000	-1.451215000

6	-5.427448000	-2.151792000	-1.062622000

1	-5.464840000	-3.025933000	-1.739796000

7	-4.173883000	-1.373443000	-1.106555000

6	-2.858280000	-1.763120000	-1.032720000

1	-2.549015000	-2.803614000	-0.943905000

7	-2.035787000	-0.706899000	-1.005564000

6	-2.872305000	0.434798000	-1.192475000

6	-2.601999000	1.835237000	-1.335868000

7	-3.739022000	2.621739000	-1.435716000

6	-5.009040000	2.110327000	-1.395001000

7	-5.254942000	0.829476000	-1.291752000

6	-4.178588000	0.001340000	-1.196973000

6	-7.223741000	-2.824732000	0.110072000

1	-7.778118000	-3.107320000	1.011539000

6	-5.707300000	-2.750115000	0.323538000

1	-5.527799000	-2.059049000	1.097984000

8	-7.521945000	-3.786914000	-0.957517000

1	-7.255726000	-4.684005000	-0.645676000

1	-5.118259000	-3.663504000	0.576739000

8	-5.771872000	0.806248000	3.319570000

1	-0.121066000	1.855462000	-1.214927000

1	-3.618704000	3.631561000	-1.531720000

1	-0.512017000	-0.692862000	-0.783449000

8	-1.495462000	2.414483000	-1.383857000

7	-6.071995000	3.073109000	-1.479448000

1	-6.867919000	2.616520000	-1.911541000

1	-6.321533000	3.396800000	-0.540291000

**G**: G _syn_ Pair**

X	Y	Z

15	-8.668356000	-1.580662000	-0.345214000

8	-7.771696000	-2.671047000	-0.863340000

8	-9.287617000	-0.844866000	-1.502837000

8	-7.828673000	-0.591775000	0.514820000

8	-7.780677000	3.166556000	-0.734531000

6	-5.099665000	1.712767000	1.312140000

6	-5.522305000	2.694199000	0.215275000

6	-6.933941000	2.207204000	-0.145729000

6	-7.440816000	1.673435000	1.209378000

6	-8.451018000	0.549003000	1.105614000

8	-6.266038000	1.244204000	1.932827000

6	-1.339550000	-0.971364000	-0.398690000

6	-2.680232000	-0.739644000	0.045107000

7	-3.701366000	-1.574485000	0.307462000

6	-4.687703000	-0.773120000	0.736912000

7	-4.340798000	0.548924000	0.769786000

6	-3.034441000	0.620148000	0.318579000

7	-2.305215000	1.698642000	0.184490000

6	-1.035074000	1.463703000	-0.305623000

1	-4.436926000	2.164228000	2.056981000

1	-5.579975000	3.699353000	0.650790000

1	-4.826446000	2.730531000	-0.623144000

1	-6.888474000	1.382770000	-0.862822000

1	-7.910618000	2.499048000	1.767434000

1	-8.857829000	0.276390000	2.057769000

1	-9.237668000	0.895324000	0.467635000

1	-5.679659000	-1.104186000	1.010585000

8	-9.757484000	-2.184739000	0.500895000

1	-7.838953000	3.932000000	-0.144964000

7	-0.601572000	0.140248000	-0.568382000

15	6.604602000	-0.761812000	2.195731000

8	8.066132000	-0.911626000	2.522832000

8	5.975885000	-2.122938000	2.046485000

8	6.457609000	0.025795000	0.860239000

6	7.625940000	0.449408000	0.151965000

1	7.938918000	-0.334192000	-0.505692000

1	8.408586000	0.661600000	0.852322000

6	7.305050000	1.664637000	-0.702999000

1	8.234237000	2.016944000	-1.162246000

8	6.409760000	1.314543000	-1.787596000

6	5.135057000	1.836753000	-1.574683000

1	4.880332000	2.530946000	-2.385338000

7	4.096015000	0.766457000	-1.642488000

6	2.756499000	1.081398000	-1.649996000

1	2.395814000	2.092507000	-1.774387000

7	1.979080000	0.044406000	-1.446127000

6	2.842596000	-1.013444000	-1.285127000

6	2.558817000	-2.352806000	-0.923612000

7	3.722094000	-3.095291000	-0.750251000

6	5.004139000	-2.602709000	-0.911586000

7	5.264916000	-1.372573000	-1.287078000

6	4.173651000	-0.595177000	-1.409139000

6	6.624613000	2.825943000	0.032271000

1	6.880015000	2.832259000	1.099212000

6	5.137653000	2.529372000	-0.194625000

1	4.777791000	1.838085000	0.569441000

8	7.042719000	4.024885000	-0.613378000

1	6.677109000	4.772385000	-0.122380000

1	4.522836000	3.433550000	-0.182577000

8	5.918070000	-0.015127000	3.308185000

8	-0.899557000	-2.158979000	-0.610538000

1	0.135069000	-2.325152000	-0.736941000

7	-0.177494000	2.397608000	-0.556563000

1	-0.611074000	3.293360000	-0.324285000

8	1.449418000	-2.903086000	-0.739091000

7	6.020680000	-3.496281000	-0.706426000

1	3.585553000	-4.065825000	-0.491223000

1	6.920073000	-3.031077000	-0.716334000

1	5.921608000	-4.066430000	0.126838000

1	0.393205000	0.076446000	-0.947765000

**G**: A _syn_ pair**

X	Y	Z

15	8.726126000	1.429274000	-0.468454000

8	7.922087000	2.600783000	-0.963770000

8	9.323613000	0.700766000	-1.642819000

8	7.805538000	0.455657000	0.331824000

8	7.576371000	-3.155864000	-1.159806000

6	5.005337000	-1.805382000	1.097299000

6	5.362552000	-2.669180000	-0.118050000

6	6.789589000	-2.213781000	-0.467867000

6	7.350363000	-1.823677000	0.915310000

6	8.390921000	-0.719222000	0.901003000

8	6.211931000	-1.415524000	1.704962000

6	1.252357000	1.141179000	0.095271000

6	2.584806000	0.818900000	0.332753000

7	3.698261000	1.597971000	0.460403000

6	4.667187000	0.729052000	0.703774000

7	4.234659000	-0.584759000	0.742053000

6	2.875356000	-0.562876000	0.502237000

7	2.040228000	-1.575016000	0.442504000

6	0.729679000	-1.233377000	0.172054000

1	4.376170000	-2.337394000	1.817988000

1	5.380392000	-3.721064000	0.192552000

1	4.647341000	-2.574005000	-0.936084000

1	6.768394000	-1.328689000	-1.108857000

1	7.803213000	-2.711261000	1.385735000

1	8.740342000	-0.497137000	1.886800000

1	9.209966000	-1.028158000	0.288094000

1	5.710112000	0.976878000	0.844013000

8	9.830043000	1.923643000	0.427793000

1	7.604922000	-3.973854000	-0.643230000

7	0.371777000	0.141413000	0.021948000

15	-6.604780000	0.621440000	2.008721000

8	-8.040857000	0.860436000	2.385073000

8	-5.876914000	1.937710000	1.941843000

8	-6.537386000	-0.073688000	0.618860000

6	-7.727174000	-0.407934000	-0.103256000

1	-8.015142000	0.413574000	-0.722694000

1	-8.519815000	-0.628707000	0.584019000

6	-7.404051000	-1.592012000	-1.000975000

1	-8.301943000	-1.881728000	-1.554214000

8	-6.411067000	-1.228639000	-1.988015000

6	-5.141292000	-1.626026000	-1.585523000

1	-4.652652000	-2.123346000	-2.430371000

7	-4.264909000	-0.444736000	-1.269492000

6	-2.977734000	-0.588642000	-0.857393000

1	-2.485875000	-1.546956000	-0.742670000

7	-2.361843000	0.560575000	-0.594985000

6	-3.309775000	1.515298000	-0.842900000

6	-3.294137000	2.932360000	-0.751321000

7	-4.403968000	3.618620000	-1.077700000

6	-5.473560000	2.927505000	-1.473204000

7	-5.615134000	1.594669000	-1.618073000

6	-4.519709000	0.926462000	-1.278929000

6	-6.812649000	-2.812891000	-0.284253000

1	-7.159634000	-2.876889000	0.754954000

6	-5.296031000	-2.551732000	-0.352720000

1	-4.959086000	-2.044760000	0.552219000

8	-7.204656000	-3.958394000	-1.032288000

1	-6.895427000	-4.743421000	-0.561710000

1	-4.736629000	-3.483288000	-0.473599000

8	-5.953782000	-0.259359000	3.040278000

8	0.797588000	2.386552000	-0.058768000

7	-0.239670000	-2.075967000	0.029829000

1	0.135711000	-3.015454000	0.165165000

7	-2.208613000	3.625285000	-0.362928000

1	-2.295096000	4.627666000	-0.283793000

1	-1.347513000	3.168119000	-0.102855000

1	-6.349085000	3.520027000	-1.724756000

1	-0.642131000	0.351650000	-0.173815000

1	1.555469000	2.994334000	0.006977000
