## Supplementary File 2s for "Higher frequency of transition mutation over transversion mutation in genomes: evidence from the binding energy calculation of base pairs using DFT"

| **A:T** | **G:C** | **A*:C** | **A:C***  **Frequency values of Base pairs** | **G:T*** | **G*:T** | **A*:A _syn_** | **A*:G _syn_** | **G**:G _syn_** | **G**: A _syn_** |
| --- | --- | --- | --- | --- | --- | --- | --- | --- | --- |
| 11 | 12 | 12 | 11 | 12 | 11 | 10 | 10 | 12 | 10 |
| 25 | 26 | 25 | 26 | 24 | 22 | 18 | 23 | 23 | 18 |
| 27 | 29 | 31 | 28 | 29 | 26 | 27 | 26 | 30 | 27 |
| 29 | 31 | 35 | 31 | 30 | 28 | 32 | 32 | 33 | 29 |
| 42 | 47 | 46 | 43 | 44 | 44 | 38 | 40 | 40 | 38 |
| 46 | 49 | 49 | 49 | 49 | 48 | 45 | 43 | 47 | 46 |
| 52 | 55 | 57 | 54 | 52 | 50 | 46 | 52 | 49 | 48 |
| 62 | 64 | 64 | 63 | 62 | 62 | 58 | 57 | 56 | 59 |
| 64 | 67 | 68 | 65 | 66 | 64 | 64 | 64 | 67 | 65 |
| 72 | 75 | 75 | 76 | 76 | 78 | 76 | 72 | 82 | 71 |
| 78 | 87 | 84 | 81 | 85 | 80 | 83 | 80 | 83 | 78 |
| 97 | 105 | 106 | 107 | 102 | 102 | 94 | 91 | 102 | 83 |
| 103 | 111 | 117 | 119 | 102 | 107 | 103 | 99 | 114 | 95 |
| 115 | 114 | 120 | 124 | 118 | 122 | 116 | 107 | 119 | 105 |
| 133 | 145 | 136 | 140 | 137 | 137 | 130 | 126 | 137 | 122 |
| 137 | 153 | 164 | 164 | 143 | 145 | 145 | 135 | 140 | 142 |
| 149 | 167 | 167 | 166 | 154 | 149 | 165 | 151 | 145 | 148 |
| 159 | 172 | 170 | 170 | 159 | 156 | 170 | 166 | 159 | 167 |
| 165 | 181 | 186 | 177 | 169 | 171 | 182 | 178 | 172 | 171 |
| 176 | 192 | 192 | 186 | 175 | 174 | 207 | 183 | 176 | 202 |
| 184 | 207 | 218 | 203 | 180 | 184 | 231 | 206 | 202 | 210 |
| 192 | 223 | 237 | 237 | 192 | 189 | 233 | 227 | 212 | 233 |
| 236 | 232 | 244 | 242 | 224 | 214 | 243 | 244 | 226 | 242 |
| 239 | 241 | 257 | 261 | 232 | 232 | 270 | 254 | 242 | 248 |
| 242 | 261 | 274 | 280 | 238 | 236 | 272 | 266 | 245 | 271 |
| 269 | 293 | 304 | 308 | 243 | 241 | 280 | 270 | 261 | 279 |
| 281 | 313 | 309 | 309 | 274 | 273 | 300 | 270 | 272 | 286 |
| 306 | 317 | 325 | 325 | 294 | 296 | 315 | 292 | 291 | 287 |
| 311 | 325 | 325 | 327 | 308 | 311 | 320 | 305 | 303 | 303 |
| 323 | 332 | 350 | 335 | 317 | 318 | 324 | 308 | 305 | 314 |
| 326 | 360 | 387 | 387 | 325 | 324 | 342 | 320 | 311 | 318 |
| 330 | 385 | 396 | 395 | 333 | 332 | 345 | 323 | 323 | 334 |
| 334 | 392 | 413 | 414 | 338 | 336 | 356 | 341 | 330 | 340 |
| 395 | 408 | 419 | 420 | 364 | 359 | 395 | 349 | 342 | 345 |
| 396 | 414 | 419 | 424 | 384 | 396 | 416 | 362 | 348 | 355 |
| 415 | 420 | 424 | 431 | 399 | 396 | 418 | 389 | 380 | 377 |
| 420 | 424 | 449 | 474 | 402 | 413 | 458 | 395 | 383 | 408 |
| 422 | 427 | 475 | 488 | 420 | 419 | 472 | 420 | 398 | 412 |
| 432 | 452 | 497 | 522 | 422 | 429 | 516 | 438 | 413 | 429 |
| 474 | 477 | 516 | 530 | 441 | 433 | 522 | 466 | 430 | 460 |
| 483 | 488 | 523 | 538 | 447 | 438 | 531 | 467 | 447 | 473 |
| 501 | 496 | 543 | 568 | 475 | 479 | 550 | 500 | 466 | 491 |
| 529 | 537 | 576 | 580 | 490 | 490 | 559 | 514 | 471 | 522 |
| 538 | 557 | 583 | 586 | 502 | 495 | 579 | 522 | 487 | 529 |
| 544 | 558 | 596 | 593 | 514 | 507 | 583 | 527 | 506 | 536 |
| 558 | 588 | 602 | 606 | 540 | 551 | 605 | 588 | 536 | 553 |
| 581 | 593 | 624 | 625 | 557 | 553 | 615 | 589 | 569 | 564 |
| 597 | 605 | 630 | 631 | 563 | 576 | 630 | 604 | 592 | 568 |
| 600 | 605 | 643 | 641 | 594 | 600 | 636 | 615 | 600 | 595 |
| 626 | 643 | 656 | 675 | 601 | 606 | 642 | 617 | 625 | 609 |
| 639 | 661 | 694 | 693 | 607 | 607 | 662 | 620 | 625 | 620 |
| 650 | 673 | 706 | 703 | 664 | 660 | 684 | 638 | 654 | 647 |
| 675 | 678 | 730 | 727 | 664 | 661 | 696 | 642 | 657 | 651 |
| 694 | 705 | 743 | 736 | 676 | 675 | 711 | 655 | 662 | 671 |
| 701 | 710 | 751 | 738 | 679 | 680 | 733 | 686 | 673 | 678 |
| 737 | 725 | 767 | 748 | 705 | 704 | 747 | 694 | 680 | 680 |
| 743 | 740 | 775 | 767 | 714 | 721 | 760 | 704 | 698 | 706 |
| 746 | 748 | 778 | 783 | 724 | 734 | 769 | 707 | 707 | 707 |
| 758 | 751 | 783 | 791 | 746 | 744 | 779 | 731 | 709 | 708 |
| 780 | 767 | 790 | 801 | 753 | 754 | 789 | 748 | 712 | 746 |
| 782 | 775 | 797 | 806 | 764 | 760 | 798 | 771 | 715 | 751 |
| 796 | 778 | 828 | 829 | 773 | 768 | 804 | 774 | 762 | 762 |
| 801 | 801 | 829 | 831 | 774 | 780 | 826 | 777 | 763 | 771 |
| 825 | 803 | 829 | 897 | 790 | 789 | 829 | 792 | 773 | 801 |
| 829 | 827 | 901 | 912 | 825 | 790 | 862 | 804 | 777 | 802 |
| 873 | 830 | 926 | 930 | 828 | 826 | 868 | 829 | 783 | 811 |
| 913 | 848 | 933 | 935 | 836 | 829 | 901 | 835 | 807 | 827 |
| 928 | 875 | 934 | 936 | 849 | 857 | 903 | 848 | 828 | 846 |
| 935 | 925 | 945 | 947 | 858 | 881 | 905 | 854 | 836 | 866 |
| 938 | 933 | 949 | 948 | 886 | 926 | 907 | 862 | 849 | 900 |
| 948 | 934 | 988 | 959 | 922 | 932 | 932 | 890 | 856 | 904 |
| 958 | 944 | 996 | 994 | 922 | 936 | 935 | 900 | 863 | 922 |
| 967 | 955 | 1003 | 1002 | 937 | 940 | 942 | 906 | 871 | 930 |
| 973 | 1000 | 1013 | 1008 | 944 | 948 | 952 | 929 | 908 | 932 |
| 992 | 1005 | 1024 | 1015 | 963 | 963 | 971 | 937 | 916 | 939 |
| 1008 | 1005 | 1031 | 1026 | 979 | 975 | 977 | 949 | 935 | 948 |
| 1011 | 1016 | 1033 | 1033 | 997 | 1008 | 980 | 954 | 944 | 973 |
| 1032 | 1028 | 1035 | 1033 | 1013 | 1011 | 986 | 970 | 950 | 973 |
| 1033 | 1032 | 1070 | 1038 | 1017 | 1017 | 1017 | 983 | 974 | 979 |
| 1035 | 1035 | 1077 | 1079 | 1032 | 1034 | 1023 | 1002 | 984 | 983 |
| 1041 | 1039 | 1089 | 1081 | 1035 | 1037 | 1035 | 1019 | 1001 | 994 |
| 1056 | 1067 | 1092 | 1089 | 1038 | 1041 | 1039 | 1022 | 1019 | 1010 |
| 1073 | 1076 | 1096 | 1098 | 1043 | 1048 | 1058 | 1025 | 1024 | 1013 |
| 1082 | 1090 | 1111 | 1113 | 1069 | 1066 | 1068 | 1035 | 1031 | 1031 |
| 1089 | 1093 | 1114 | 1117 | 1074 | 1074 | 1093 | 1065 | 1048 | 1047 |
| 1106 | 1107 | 1132 | 1141 | 1084 | 1089 | 1096 | 1073 | 1050 | 1052 |
| 1113 | 1116 | 1142 | 1146 | 1090 | 1094 | 1100 | 1077 | 1067 | 1068 |
| 1117 | 1122 | 1143 | 1171 | 1093 | 1100 | 1118 | 1091 | 1079 | 1089 |
| 1141 | 1143 | 1170 | 1175 | 1108 | 1115 | 1144 | 1093 | 1091 | 1095 |
| 1146 | 1145 | 1172 | 1185 | 1118 | 1118 | 1148 | 1113 | 1094 | 1111 |
| 1176 | 1151 | 1180 | 1202 | 1121 | 1131 | 1181 | 1130 | 1096 | 1121 |
| 1182 | 1175 | 1214 | 1212 | 1133 | 1147 | 1187 | 1138 | 1115 | 1142 |
| 1186 | 1182 | 1231 | 1232 | 1147 | 1154 | 1190 | 1140 | 1133 | 1144 |
| 1219 | 1196 | 1237 | 1237 | 1164 | 1182 | 1221 | 1164 | 1140 | 1181 |
| 1232 | 1215 | 1242 | 1242 | 1182 | 1183 | 1226 | 1192 | 1145 | 1189 |
| 1237 | 1223 | 1254 | 1278 | 1186 | 1210 | 1235 | 1206 | 1150 | 1218 |
| 1242 | 1228 | 1278 | 1287 | 1205 | 1224 | 1241 | 1215 | 1185 | 1222 |
| 1254 | 1238 | 1292 | 1293 | 1220 | 1228 | 1244 | 1222 | 1205 | 1224 |
| 1281 | 1240 | 1296 | 1297 | 1228 | 1238 | 1264 | 1226 | 1209 | 1232 |
| 1293 | 1278 | 1315 | 1314 | 1232 | 1239 | 1275 | 1226 | 1217 | 1234 |
| 1297 | 1295 | 1316 | 1320 | 1237 | 1262 | 1292 | 1238 | 1223 | 1239 |
| 1313 | 1296 | 1330 | 1333 | 1241 | 1294 | 1296 | 1260 | 1225 | 1259 |
| 1320 | 1321 | 1333 | 1336 | 1265 | 1296 | 1311 | 1263 | 1238 | 1262 |
| 1333 | 1324 | 1338 | 1337 | 1289 | 1301 | 1321 | 1279 | 1239 | 1289 |
| 1336 | 1334 | 1338 | 1340 | 1295 | 1318 | 1328 | 1292 | 1273 | 1297 |
| 1337 | 1336 | 1344 | 1349 | 1306 | 1336 | 1337 | 1299 | 1291 | 1311 |
| 1340 | 1339 | 1347 | 1367 | 1318 | 1337 | 1338 | 1317 | 1293 | 1312 |
| 1349 | 1343 | 1365 | 1369 | 1324 | 1340 | 1340 | 1326 | 1304 | 1331 |
| 1358 | 1367 | 1368 | 1373 | 1332 | 1344 | 1350 | 1334 | 1319 | 1334 |
| 1368 | 1368 | 1394 | 1395 | 1336 | 1348 | 1352 | 1337 | 1328 | 1336 |
| 1370 | 1381 | 1396 | 1396 | 1338 | 1370 | 1355 | 1340 | 1335 | 1338 |
| 1386 | 1395 | 1403 | 1405 | 1341 | 1372 | 1366 | 1343 | 1336 | 1341 |
| 1395 | 1395 | 1407 | 1417 | 1347 | 1378 | 1370 | 1347 | 1340 | 1347 |
| 1396 | 1407 | 1440 | 1424 | 1368 | 1394 | 1376 | 1355 | 1348 | 1356 |
| 1403 | 1415 | 1446 | 1446 | 1369 | 1397 | 1387 | 1357 | 1354 | 1366 |
| 1422 | 1446 | 1447 | 1448 | 1379 | 1398 | 1399 | 1371 | 1360 | 1368 |
| 1427 | 1448 | 1448 | 1451 | 1394 | 1400 | 1401 | 1373 | 1363 | 1372 |
| 1433 | 1451 | 1453 | 1453 | 1397 | 1413 | 1418 | 1387 | 1371 | 1372 |
| 1446 | 1455 | 1489 | 1483 | 1404 | 1438 | 1420 | 1390 | 1377 | 1393 |
| 1448 | 1478 | 1498 | 1495 | 1413 | 1446 | 1423 | 1393 | 1380 | 1394 |
| 1451 | 1493 | 1500 | 1498 | 1439 | 1447 | 1436 | 1414 | 1393 | 1402 |
| 1470 | 1498 | 1525 | 1500 | 1446 | 1448 | 1442 | 1416 | 1396 | 1413 |
| 1484 | 1499 | 1558 | 1528 | 1449 | 1451 | 1442 | 1432 | 1409 | 1423 |
| 1490 | 1527 | 1573 | 1535 | 1451 | 1480 | 1446 | 1439 | 1422 | 1440 |
| 1497 | 1550 | 1578 | 1606 | 1458 | 1490 | 1481 | 1442 | 1434 | 1441 |
| 1500 | 1575 | 1601 | 1615 | 1477 | 1497 | 1497 | 1458 | 1439 | 1443 |
| 1506 | 1603 | 1654 | 1626 | 1489 | 1498 | 1504 | 1481 | 1441 | 1468 |
| 1524 | 1645 | 1683 | 1668 | 1494 | 1503 | 1507 | 1486 | 1443 | 1483 |
| 1533 | 1668 | 1709 | 1713 | 1498 | 1509 | 1534 | 1501 | 1480 | 1497 |
| 1607 | 1698 | 1740 | 1794 | 1499 | 1530 | 1568 | 1510 | 1485 | 1500 |
| 1615 | 1711 | 2714 | 2938 | 1505 | 1536 | 1579 | 1550 | 1498 | 1510 |
| 1630 | 1721 | 2985 | 2985 | 1524 | 1553 | 1603 | 1558 | 1503 | 1516 |
| 1692 | 1773 | 2986 | 2990 | 1552 | 1607 | 1609 | 1580 | 1524 | 1553 |
| 1725 | 2872 | 3054 | 3055 | 1585 | 1638 | 1628 | 1589 | 1550 | 1600 |
| 1783 | 2984 | 3056 | 3056 | 1602 | 1666 | 1686 | 1594 | 1569 | 1610 |
| 2986 | 2986 | 3090 | 3090 | 1633 | 1699 | 1705 | 1615 | 1592 | 1621 |
| 2986 | 2988 | 3108 | 3108 | 1675 | 1717 | 2976 | 1670 | 1598 | 1666 |
| 3034 | 3052 | 3108 | 3110 | 1699 | 1757 | 3050 | 1680 | 1633 | 1673 |
| 3055 | 3056 | 3128 | 3116 | 1713 | 2757 | 3054 | 1727 | 1649 | 1690 |
| 3057 | 3084 | 3143 | 3131 | 1765 | 2977 | 3066 | 3010 | 1671 | 2832 |
| 3081 | 3109 | 3145 | 3145 | 2825 | 2985 | 3076 | 3039 | 1694 | 2998 |
| 3088 | 3111 | 3215 | 3149 | 2961 | 3052 | 3085 | 3054 | 1738 | 3048 |
| 3099 | 3123 | 3226 | 3201 | 2981 | 3054 | 3088 | 3055 | 2237 | 3055 |
| 3105 | 3143 | 3240 | 3226 | 2987 | 3056 | 3107 | 3084 | 2602 | 3063 |
| 3111 | 3148 | 3257 | 3246 | 3052 | 3083 | 3108 | 3086 | 3000 | 3080 |
| 3125 | 3232 | 3262 | 3261 | 3054 | 3106 | 3148 | 3087 | 3041 | 3088 |
| 3142 | 3242 | 3503 | 3529 | 3056 | 3111 | 3153 | 3091 | 3054 | 3105 |
| 3145 | 3254 | 3696 | 3647 | 3079 | 3112 | 3191 | 3115 | 3056 | 3112 |
| 3147 | 3425 | 3788 | 3788 | 3083 | 3113 | 3197 | 3152 | 3083 | 3149 |
| 3200 | 3670 | 3788 | 3789 | 3107 | 3122 | 3207 | 3158 | 3086 | 3155 |
| 3217 | 3705 |  |  | 3111 | 3144 | 3262 | 3212 | 3091 | 3198 |
| 3264 | 3788 |  |  | 3114 | 3144 | 3275 | 3258 | 3107 | 3248 |
| 3276 | 3789 |  |  | 3125 | 3147 | 3512 | 3316 | 3151 | 3280 |
| 3666 |  |  |  | 3143 | 3225 | 3643 | 3474 | 3156 | 3518 |
| 3788 |  |  |  | 3146 | 3257 | 3787 | 3566 | 3279 | 3586 |
| 3789 |  |  |  | 3146 | 3279 | 3817 | 3600 | 3287 | 3729 |
|  |  |  |  | 3225 | 3683 |  | 3672 | 3506 | 3732 |
|  |  |  |  | 3257 | 3788 |  | 3787 | 3557 | 3787 |
|  |  |  |  | 3283 | 3788 |  | 3816 | 3600 | 3816 |
|  |  |  |  | 3676 |  |  |  | 3660 |  |
|  |  |  |  | 3788 |  |  |  | 3787 |  |
|  |  |  |  | 3788 |  |  |  | 3816 |  |

**Frequency values of Single Base**

| A | A* | T | T* | G | G* | C | C* | G** | A _syn_ | G _syn_ |
| --- | --- | --- | --- | --- | --- | --- | --- | --- | --- | --- |
| 6 | 5 | 11 | 11 | 5 | 5 | 6 | 9 | 4 | 14 | 7 |
| 15 | 15 | 15 | 16 | 12 | 12 | 19 | 16 | 14 | 23 | 18 |
| 18 | 22 | 39 | 37 | 16 | 15 | 31 | 30 | 20 | 40 | 34 |
| 37 | 36 | 42 | 41 | 33 | 36 | 45 | 45 | 34 | 52 | 39 |
| 57 | 58 | 63 | 64 | 56 | 56 | 62 | 62 | 61 | 54 | 69 |
| 63 | 64 | 92 | 97 | 60 | 62 | 70 | 71 | 63 | 84 | 82 |
| 105 | 99 | 112 | 110 | 91 | 92 | 105 | 108 | 91 | 105 | 107 |
| 109 | 110 | 146 | 147 | 104 | 103 | 156 | 156 | 99 | 140 | 129 |
| 164 | 153 | 153 | 164 | 143 | 141 | 167 | 159 | 135 | 170 | 138 |
| 181 | 173 | 171 | 179 | 163 | 171 | 177 | 170 | 169 | 206 | 168 |
| 227 | 219 | 190 | 226 | 170 | 205 | 190 | 202 | 196 | 241 | 199 |
| 241 | 242 | 233 | 232 | 208 | 216 | 218 | 266 | 226 | 266 | 223 |
| 273 | 260 | 270 | 274 | 221 | 239 | 266 | 309 | 244 | 271 | 242 |
| 298 | 303 | 280 | 279 | 241 | 241 | 316 | 328 | 280 | 285 | 295 |
| 310 | 324 | 310 | 308 | 292 | 291 | 325 | 383 | 297 | 321 | 303 |
| 328 | 336 | 329 | 328 | 314 | 313 | 369 | 389 | 312 | 338 | 321 |
| 343 | 392 | 338 | 338 | 336 | 322 | 387 | 412 | 333 | 361 | 349 |
| 394 | 421 | 391 | 386 | 338 | 333 | 417 | 423 | 372 | 399 | 360 |
| 420 | 471 | 403 | 401 | 357 | 359 | 425 | 479 | 408 | 454 | 404 |
| 471 | 511 | 417 | 420 | 398 | 406 | 482 | 520 | 426 | 470 | 447 |
| 525 | 532 | 424 | 437 | 422 | 423 | 517 | 558 | 474 | 508 | 462 |
| 541 | 579 | 470 | 477 | 459 | 478 | 543 | 594 | 484 | 522 | 467 |
| 575 | 583 | 488 | 505 | 475 | 497 | 582 | 623 | 545 | 547 | 488 |
| 579 | 618 | 545 | 550 | 484 | 560 | 593 | 672 | 569 | 566 | 524 |
| 584 | 627 | 601 | 601 | 553 | 572 | 622 | 680 | 596 | 602 | 594 |
| 627 | 634 | 644 | 609 | 591 | 575 | 687 | 729 | 623 | 634 | 612 |
| 635 | 656 | 685 | 647 | 610 | 599 | 728 | 743 | 642 | 677 | 632 |
| 682 | 687 | 689 | 677 | 629 | 624 | 744 | 757 | 664 | 706 | 666 |
| 698 | 725 | 741 | 730 | 658 | 649 | 770 | 775 | 667 | 746 | 688 |
| 737 | 771 | 750 | 746 | 672 | 674 | 788 | 788 | 674 | 757 | 694 |
| 785 | 778 | 762 | 768 | 688 | 679 | 791 | 812 | 700 | 794 | 719 |
| 807 | 802 | 772 | 782 | 696 | 698 | 828 | 829 | 720 | 801 | 761 |
| 829 | 830 | 826 | 825 | 721 | 734 | 938 | 936 | 749 | 867 | 773 |
| 904 | 904 | 856 | 863 | 766 | 771 | 948 | 957 | 771 | 885 | 813 |
| 928 | 912 | 933 | 933 | 775 | 792 | 968 | 981 | 818 | 893 | 847 |
| 937 | 929 | 948 | 948 | 828 | 828 | 1010 | 1007 | 828 | 916 | 861 |
| 949 | 936 | 955 | 960 | 850 | 849 | 1021 | 1017 | 846 | 922 | 862 |
| 970 | 949 | 998 | 998 | 905 | 919 | 1035 | 1032 | 934 | 954 | 874 |
| 1001 | 1002 | 1021 | 1013 | 924 | 937 | 1062 | 1076 | 936 | 979 | 904 |
| 1016 | 1021 | 1031 | 1035 | 948 | 948 | 1073 | 1093 | 948 | 981 | 942 |
| 1034 | 1034 | 1040 | 1041 | 1000 | 1002 | 1094 | 1117 | 963 | 1009 | 977 |
| 1058 | 1070 | 1062 | 1050 | 1029 | 1024 | 1119 | 1145 | 1007 | 1038 | 985 |
| 1089 | 1091 | 1099 | 1093 | 1032 | 1035 | 1147 | 1155 | 1031 | 1066 | 1028 |
| 1114 | 1115 | 1118 | 1119 | 1059 | 1056 | 1175 | 1178 | 1047 | 1093 | 1056 |
| 1141 | 1139 | 1143 | 1140 | 1072 | 1068 | 1205 | 1207 | 1088 | 1118 | 1063 |
| 1178 | 1161 | 1175 | 1154 | 1088 | 1090 | 1236 | 1236 | 1103 | 1173 | 1097 |
| 1192 | 1180 | 1188 | 1190 | 1096 | 1115 | 1264 | 1281 | 1116 | 1215 | 1111 |
| 1233 | 1207 | 1235 | 1235 | 1129 | 1143 | 1289 | 1292 | 1143 | 1221 | 1119 |
| 1241 | 1235 | 1253 | 1261 | 1132 | 1181 | 1313 | 1304 | 1181 | 1230 | 1145 |
| 1266 | 1238 | 1294 | 1279 | 1164 | 1209 | 1327 | 1329 | 1211 | 1258 | 1208 |
| 1296 | 1246 | 1312 | 1295 | 1183 | 1233 | 1334 | 1335 | 1230 | 1291 | 1226 |
| 1318 | 1297 | 1332 | 1317 | 1222 | 1242 | 1365 | 1366 | 1235 | 1315 | 1233 |
| 1336 | 1332 | 1337 | 1334 | 1236 | 1250 | 1380 | 1396 | 1245 | 1322 | 1249 |
| 1340 | 1339 | 1366 | 1336 | 1243 | 1296 | 1398 | 1397 | 1295 | 1334 | 1300 |
| 1347 | 1343 | 1370 | 1359 | 1289 | 1322 | 1435 | 1418 | 1301 | 1345 | 1319 |
| 1369 | 1369 | 1394 | 1368 | 1307 | 1336 | 1444 | 1436 | 1322 | 1358 | 1336 |
| 1385 | 1376 | 1395 | 1394 | 1332 | 1340 | 1500 | 1444 | 1333 | 1374 | 1346 |
| 1395 | 1395 | 1405 | 1402 | 1337 | 1364 | 1504 | 1498 | 1339 | 1390 | 1353 |
| 1400 | 1414 | 1414 | 1442 | 1342 | 1370 | 1557 | 1508 | 1353 | 1391 | 1370 |
| 1415 | 1430 | 1443 | 1449 | 1355 | 1378 | 1628 | 1608 | 1369 | 1402 | 1380 |
| 1446 | 1442 | 1448 | 1459 | 1370 | 1395 | 1660 | 1644 | 1389 | 1427 | 1393 |
| 1446 | 1448 | 1467 | 1480 | 1396 | 1406 | 1726 | 1809 | 1395 | 1441 | 1409 |
| 1492 | 1474 | 1494 | 1503 | 1404 | 1446 | 2989 | 2998 | 1403 | 1484 | 1432 |
| 1500 | 1498 | 1503 | 1524 | 1411 | 1448 | 3056 | 3057 | 1442 | 1503 | 1439 |
| 1513 | 1524 | 1615 | 1546 | 1447 | 1486 | 3098 | 3104 | 1447 | 1519 | 1490 |
| 1606 | 1568 | 1770 | 1656 | 1448 | 1499 | 3123 | 3109 | 1497 | 1596 | 1495 |
| 1619 | 1615 | 1799 | 1747 | 1495 | 1519 | 3163 | 3160 | 1512 | 1629 | 1542 |
| 1664 | 1699 | 2838 | 2837 | 1501 | 1523 | 3232 | 3240 | 1552 | 1671 | 1598 |
| 2983 | 2988 | 3015 | 3014 | 1571 | 1617 | 3241 | 3251 | 1642 | 3040 | 1602 |
| 3053 | 3053 | 3057 | 3056 | 1579 | 1649 | 3618 | 3510 | 1645 | 3078 | 1696 |
| 3090 | 3091 | 3067 | 3072 | 1602 | 1662 | 3758 | 3613 | 1690 | 3086 | 1804 |
| 3107 | 3110 | 3104 | 3103 | 1670 | 2980 | 3789 | 3788 | 2994 | 3108 | 3022 |
| 3146 | 3145 | 3111 | 3117 | 1813 | 3051 |  |  | 3054 | 3150 | 3027 |
| 3195 | 3204 | 3163 | 3165 | 2964 | 3085 |  |  | 3086 | 3194 | 3070 |
| 3273 | 3280 | 3172 | 3177 | 2981 | 3109 |  |  | 3109 | 3274 | 3076 |
| 3600 | 3470 | 3279 | 3281 | 3051 | 3144 |  |  | 3147 | 3588 | 3092 |
| 3742 | 3605 | 3608 | 3745 | 3088 | 3267 |  |  | 3276 | 3725 | 3166 |
| 3788 | 3787 | 3788 | 3789 | 3109 | 3628 |  |  | 3527 | 3816 | 3264 |
|  |  |  |  | 3143 | 3751 |  |  | 3603 |  | 3594 |
|  |  |  |  | 3269 | 3773 |  |  | 3740 |  | 3663 |
|  |  |  |  | 3597 | 3788 |  |  | 3787 |  | 3815 |
|  |  |  |  | 3617 |  |  |  |  |  |  |
|  |  |  |  | 3747 |  |  |  |  |  |  |
|  |  |  |  | 3788 |  |  |  |  |  |  |
